# A role for thalamic projection GABAergic neurons in circadian responses to light

**DOI:** 10.1101/2022.02.24.481804

**Authors:** O. Brock, C.E. Gelegen, I. Salgarella, P. Sully, P. Jager, L. Menage, I. Mehta, J. Jęczmień-Łazur, D. Djama, L. Strother, A. Coculla, A. Vernon, S. Brickley, P. Holland, S. Cooke, A. Delogu

**Author notes:** equal contribution (C.E. Gelegen, I. Salgarella, P. Sully).

## Abstract

The thalamus is an important hub for sensory information and participates in sensory perception, regulation of attention, arousal and sleep. These functions are executed primarily by glutamatergic thalamocortical neurons that extend axons to the cortex and initiate cortico-thalamocortical connectional loops. However, the thalamus also contains projection GABAergic neurons that do not engage in direct communication with the cortex. Here, we have harnessed recent insight into the development of the intergeniculate (IGL), the ventrolateral geniculate (LGv) and the perihabenula (pHB) to specifically target and manipulate thalamic projection GABAergic neurons in female and male mice. Our results show that thalamic GABAergic neurons of the IGL and LGv receive retinal input from diverse classes of ipRGCs, but not from the M1 ipRGC type, while those in the pHB lack direct retinal input. We describe the synergistic role of the photoreceptor melanopsin and the thalamic neurons of the IGL/LGv in circadian entrainment to dim light. We identify a requirement for the thalamic IGL/LGv in the rapid changes in vigilance states associated with circadian light transitions. Furthermore, we map a previously undescribed thalamic network of developmentally related GABAergic neurons in the IGL/LGv complex and the pHB potentially involved in light-dependent mood regulation.

**Significance statement:** The intergeniculate leaflet and ventral geniculate nucleus are part of the extended circadian system and mediate some non-image-forming visual functions. Here we show that each of these structures has a thalamic (dorsal) as well as prethalamic (ventral) developmental origin. We map the retinal input to thalamus-derived cells in the IGL/LGv complex and discover that while ipRGC input is dominant, this is not likely to originate from M1-ipRGCs. We describe the extent of similarity in synaptic input to developmentally related cells in the IGL/LGv and in the perihabenula nucleus (pHB). We implicate thalamic cells in the IGL/LGv in vigilance state transitions at circadian light changes and in overt behavioural entrainment to dim light, the latter exacerbated by concomitant loss of melanopsin expression.

## Introduction

GABAergic projection neurons are present at the rostroventral and dorsocaudal edges of the mouse thalamus to form the intergeniculate leaflet (IGL) (Morin and Blanchard, 1999, 2001, 2005) and perihabenula nucleus (pHB) (An et al., 2020; Fernandez et al., 2018), respectively. Contiguous with the IGL, but within the largely GABAergic prethalamic territory, is the ventral lateral geniculate nucleus (LGv) (Monavarfeshani et al., 2017; Moore et al., 2000; Morin and Blanchard, 2005; Sabbagh et al., 2020). Despite their largely distinct ontogeny (Delogu et al., 2012; Inamura et al., 2011; Jeong et al., 2011; Puelles et al., 2020; Suzuki-Hirano et al., 2011; Virolainen et al., 2012; Vue et al., 2007; Yuge et al., 2011), functional studies have often grouped the IGL and LGv together, based on GABA expression, anatomical proximity and, to some degree, shared patterns of connectivity.

The IGL and LGv are the source of the geniculohypothalamic tract (Harrington, 1997; Moore, 1989; Morin and Blanchard, 1999, 2001; Pu and Pickard, 1996) that enables regulation of the circadian clock in the suprachiasmatic nucleus (SCN) (Harrington and Rusak, 1986; Johnson et al., 1989; Lewandowski and Usarek, 2002)(Fernandez et al., 2020; Hanna et al., 2017; Huhman and Albers, 1994; Huhman et al., 1995; Huhman et al., 1996; Shibata and Moore, 1993). The geniculohypothalamic tract is believed to be the conduit for integrated photic (Harrington and Rusak, 1989; Morin and Studholme, 2014b; Zhang and Rusak, 1989) and non-photic (Janik and Mrosovsky, 1994; Johnson et al., 1988; Kuroda et al., 1997; Marchant et al., 1997; Maywood et al., 2002; Maywood et al., 1997) cues that contribute to circadian entrainment to relevant external and internal variables. Retinal input is mostly from intrinsically photosensitive retinal ganglion cells (ipRGCs) (An et al., 2020; Beier et al., 2020; Fernandez et al., 2018; Guler et al., 2008; Hattar et al., 2006; Huang et al., 2019), while non-photic cues are thought to propagate via neurons of the ascending arousal system (Blasiak and Lewandowski, 2003; Marchant et al., 1997; Meyer-Bernstein and Morin, 1996; Smith et al., 2015; Vrang et al., 2003).

Neurons in the IGL/LGv were shown to participate in mood regulation via inhibitory synapses onto lateral habenula (LH) neurons (Huang et al., 2019) and to contribute to photosomnolence in mice exposed to unexpected light at night (Shi et al., 2020). Related mood-regulatory functions are attributed to the pHB (An et al., 2020; Fernandez et al., 2018). Thalamic GABAergic projection neurons are specified during embryonic development within the rostral portion of the second diencephalic prosomere (p2) (Kataoka and Shimogori, 2008; Martinez-Ferre and Martinez, 2012; Nakagawa, 2019; Puelles, 2019; Puelles and Rubenstein, 1993, 2003; Rubenstein et al., 1994; Vue et al., 2007) and can be defined by expression of the transcription factor gene *Sox14* (Delogu et al., 2012; Sellers et al., 2014; Virolainen et al., 2012; Vue et al., 2007). Tangential cell migration during embryogenesis distributes thalamic GABAergic precursors from the prospective IGL, to the developing LGv and pHB (Delogu et al., 2012). Hence, the mature IGL, LGv and pHB are characterised by a heterogenous cellular composition that includes *Sox14^+^* GABAergic thalamic neurons. While the sparse interneurons of the mouse thalamocortical nuclei also express *Sox14*, these local circuit cells have a distinctive mesencephalic origin (Jager et al., 2021; Jager et al., 2016).

Here, we used stereotaxic injections in the *Sox14^Cre^* mouse to enable the characterisation of the thalamic component of anatomical regions with complex embryonic ontogeny. We demonstrate that circadian optogenetic stimulation of the *Sox14^+^* neurons in the IGL/LGv is sufficient to reset circadian motor activity rhythms in the absence of other light cues. Upon cell ablation, we show that the thalamic component of the IGL/LGv plays a synergistic role with melanopsin photodetection to ensure photoentrainment to dim light and participates in the regulation of vigilance state transitions. We map synaptic input to thalamic *Sox14^+^* neurons in the IGL/LGv and pHB and provide evidence of a potential new pathway for mood regulation.

## Results

### The IGL/LGv complex contains cells of thalamic as well as prethalamic origins

The IGL and the LGv are thought to arise from distinct progenitor domains in the thalamic prosomere 2 and prethalamic prosomere 3, respectively (Kataoka and Shimogori, 2008; Puelles et al., 2013; Virolainen et al., 2012; Vue et al., 2007). We and others have shown that radial migration of prosomere 2 *Sox14^+^* precursors into the thalamic mantle zone generates the IGL primordium (Delogu et al., 2012; Jeong et al., 2011; Vue et al., 2007). In the developmental window between gestational day (E) E11.5-E14.5, different subsets of *Sox14^+^* neurons display tangential migratory behaviour from this location, reaching the thalamus-epithalamus border first, at E12.5 to coalesce in the presumptive pHB nucleus (Fig. 1A) and then, in a subsequent wave of rostroventral migration, seeding the presumptive LGv with neurons of thalamic origin (Delogu et al., 2012; Jeong et al., 2011; Virolainen et al., 2012; Vue et al., 2007) (Fig. 1A,Q). To assess whether prethalamic neurons also contribute to the mature IGL (Fig. 1A), we mapped the fate of prethalamic GABAergic lineages in the IGL at postnatal day 21 (P21), using the prethalamic Cre-driver mouse line *Dlx5/6^Cre^* (Jager et al., 2021; Monory et al., 2006; Puelles et al., 2020) crossed with the conditional reporter line *Rosa26-CAG-Sun1/sfGFP* (*R26^lsl-nGFP^*) (Mo et al., 2015). To assist with the anatomical delineation of the IGL, we co-labelled coronal tissue sections of the lateral geniculate with an antibody against neuropeptide Y (Npy), a known marker for a subset of IGL neurons, and counted the proportion of neurons (NeuN^+^) within IGL boundaries that have prethalamic origin (NeuN^+^nGFP^+^; Fig. 1B). This analysis revealed that about a quarter of the neurons in the IGL are of prethalamic origin (Fig. 1C; 25.69 % ± 1.17 %, mean ± s.e.m., n = 3 mice). The calcium binding proteins calbindin (Calb1) and parvalbumin (Pvalb) mark different cell types in the mature IGL/LGv complex (Sabbagh et al., 2020). Using the *Sox14^Gfp/+^* (Delogu et al., 2012) and the *Dlx5/6^Cre^;R26^lsl-nGFP^* reporter lines to label thalamic and prethalamic IGL/LGv lineages, respectively, we noted a similar proportion of Calb1^+^ IGL/LGv neuron subsets in both developmental classes. In the IGL 61.40 % ± 5.46 % of Calb1^+^ cells belonged to the thalamic *Sox14^+^* developmental class and 23.53 % ± 1.94 % to the prethalamic *Dlx5/6^+^* class; in the LGv 8.1 % ± 5.56 % belonged to the thalamic *Sox14^+^* developmental class and 84.17 % ± 5.56 % to the prethalamic *Dlx5/6^+^* class (Fig. 1D,E,F,G; mean ± s.e.m., n = 3 mice per genotype). The Pvalb^+^ subtype was virtually absent from the IGL and found exclusively in LGv lineages of prethalamic origin (Fig. 1H,I,J,K; n = 3 mice per genotype). Hence, while each developmental class clearly differentiates further into several molecularly and functionally distinct cell types (Morin and Blanchard, 1995; Morin and Blanchard, 2001; Sabbagh et al., 2020) conventional mature cell markers may not always reflect developmental origin (e.g. Calb1). Importantly, progenitors from the thalamic and prethalamic primordium contribute to the formation of the mature IGL and LGv without clear spatial segregation of developmental lineage classes between the two anatomical regions. To visualise the pattern of axonal projections from the *Sox14^+^* IGL/LGv neurons in the mature brain, we injected a Cre-dependent adenoassociated virus (AAV) expressing a cell membrane localised GFP (Matsuda and Cepko, 2007) (AAV2/1 Ef1a-DIO-mGFP) in the IGL/LGv of Sox14*^Cre/+^* mice (Jager et al., 2016) at weaning age and imaged the brain-wide extent of GFP-labelled axons 3 weeks later (Fig. 1L). Although we did not conduct a detailed analysis of axonal projections, we noted that overall, the pattern of efferent projections of the *Sox14^+^* IGL/LGv neurons was consistent with earlier reports for the anatomically defined IGL and LGv (Moore et al., 2000; Morin and Blanchard, 1995; Morin and Blanchard, 1999, 2005).

**Figure 1.**
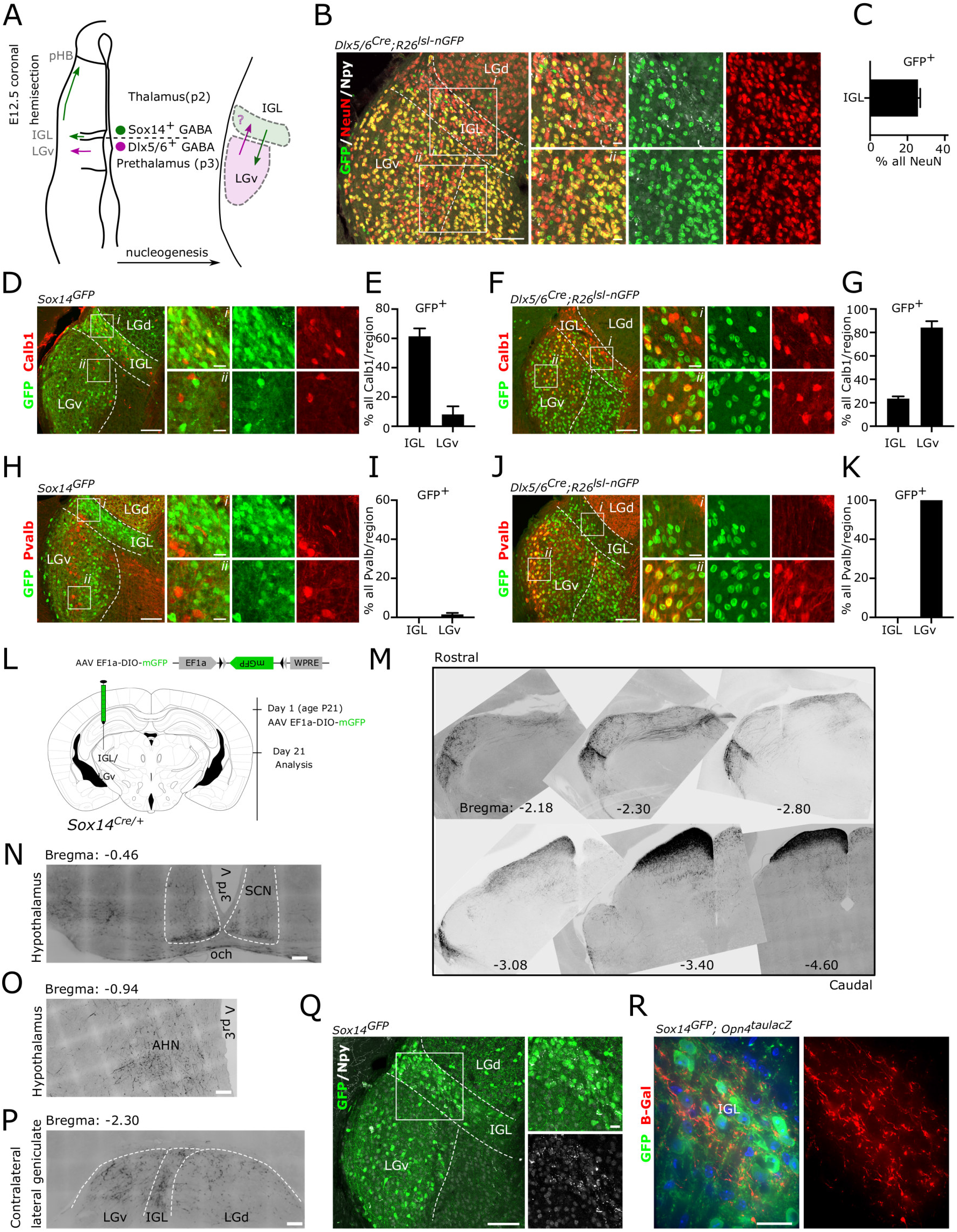
Thalamic and prethalamic lineages in the IGL/LGv complex. ***A***, schematic representation of the left diencephalon along the coronal plane at the time of peak embryonic neurogenesis (E12.5) illustrating the two main sources of GABAergic neurons for the IGL and the LGv. Radial migration of prethalamic (p3) GABAergic precursors (magenta) generates the bulk of the LGv, while radial migration of thalamic (p2) GABAergic precursors (green) generate the bulk of the IGL. Tangential migration of thalamic GABAergic precursors contributes to the formation of the pHB and to the cellular complexity of the LGv. The possibility of a complementary contribution of prethalamic GABAergic precursors to the IGL is tested using the prethalamic GABAergic driver *Dlx5/6^Cre^*. ***B***, representative image illustrating the presence of neurons (NeuN^+^) with prethalamic origins (GFP^+^) in the IGL. Note the presence of NeuN^+^GFP^neg^ neurons in the LGv, consistent with the thalamic origin of some LGv neurons and of NeuN^neg^GFP^+^ glia in the LGd. ***C***, quantification of the fraction of IGL neurons with prethalamic origin. ***D,E***, representative images and quantification of the mosaic expression of the calbindin protein (Calb1) among thalamic (*Sox14^GFP/+^*) lineages in the IGL and LGv. ***F***,***G***, representative images and quantification of the mosaic expression of Calb1 in prethalamic (*Dlx5/6^Cre^;R26^lsl-nGFP^*) lineages in the IGL and LGv. ***H***,***I***,***J***,***K***, representative images and quantification of the expression of Pvalb in thalamic and prethalamic lineages of the IGL and the LGv. ***L***, schematic illustration of timeline of the AAV injection strategy used to label axonal projections of the *Sox14^+^* IGL/LGv neurons in the adult brain. ***M***, inverted grey scale images from representative rostrocaudal levels of the IGL/LGv, demonstrating the widespread presence of dark GFP labelled fibres (see **L** for strategy) projecting away from the IGL/LGv and towards other diencephalic and mesencephalic structures. ***N***,***O***,***P***, higher magnification images from the experiment in **L** showing the presence of sparse GFP labelled fibres in the SCN, the AHN and the contralateral geniculate. ***Q***, an illustrative example of the location of thalamic *Sox14^+^* neurons in the IGL and the LGv at 3 weeks of age. ***R***, an example image showing the incoming ipRGC axons (b-gal) in the region occupied by *Sox14****^+^*** neurons in the IGL using the *Opn4^taulacZ/+^*;*Sox14^GFP/+^* double transgenic mouse. Notably, dense innervation was seen in the superficial gray and optic layers of the superior colliculus (Fig. 1M), while few immunoreactive fibres were present at the ventral edge of the SCN (Fig. 1N), sparse and diffuse in the hypothalamus (e.g. anterior hypothalamic nucleus, AHN, Fig. 1O) and in all three subdivisions of the contralateral lateral geniculate (LGN, Fig. 1P).

We had previously shown that *Sox14^+^* neurons in the IGL/LGv establish synaptic connectivity within the non-image-forming circuitry that mediates the pupillary light reflex (Delogu et al., 2012). Taking advantage of the strong GFP expression from the *Sox14^Gfp/+^* mouse reporter line in the juvenile brain (Fig. 1Q), we crossed this reporter line with the *Opn4^taulacZ/+^*, which labels mostly the M1 subtype of ipRGCs (Baver et al., 2008; Hattar et al., 2006; Hattar et al., 2002). The *Opn4^taulacZ/+^*;*Sox14^Gfp/+^* double transgenic mouse line confirmed the presence of GFP^+^ neurons within axonal projections labelled by the *taulacZ* reporter construct (Fig. 1R), consistent with several reports that propose M1 ipRGC innervation of the IGL/LGv. However, the discovery of additional prethalamic lineages in the IGL (Fig. 1B,C) raises the possibility that developmentally defined cell classes may receive selective ipRGC-subtype innervation. The *Sox14Gfp* and *Sox14Cre* mouse lines are suitable tools to resolve the developmental complexity of the IGL/LGv by restricting genetic labelling and manipulations exclusively to neurons of thalamic origin.

### Retinal input to the *Sox14^+^* IGL/LGv originates from non-M1 ipRGCs

We sought to investigate the extent and diversity of retinal input to the thalamic component of the IGL/LGv, defined by *Sox14* expression. For this purpose, we applied the modified rabies virus technology (Fig. 2A), guiding primary infection of a glycoprotein-deleted, GFP-expressing and avian-pseudotyped rabies (SADB19 ΔG-eGFP, EnvA; in short RVdG) to the *Sox14* neurons of the IGL/LGv. Target neurons were primed for RVdG infection by stereotaxic injection in *Sox14^Cre/+^* transgenic mice of equimolar amounts of two Cre-dependent AAVs (Fig. 2A) expressing the avian receptor TVA linked via a self-cleavage peptide sequence to the red fluorescent reporter mCherry (AAV2/1 Ef1a-flex-TVA-mCherry) and the codon-optimised version of the rabies glycoprotein (G) gene (AAV2/1 CAG-flex-oG). Our injection strategy reliably targeted the IGL/LGv, as indicated by the cumulative total number of primary infected neurons (GFP^+^mCherry^+^; Fig. 2B) from all thalamus-containing sections in the target region (IGL: 308, LGv: 39; n = 6 brains; Fig. 2C), while off target labelling was occasionally observed in some of the sparse *Sox14^+^* thalamic interneurons along the trajectory of the stereotaxic injection (Fig. 2C; LGd, LP, NOT: 13; n = 6 mice).

**Figure 2.**
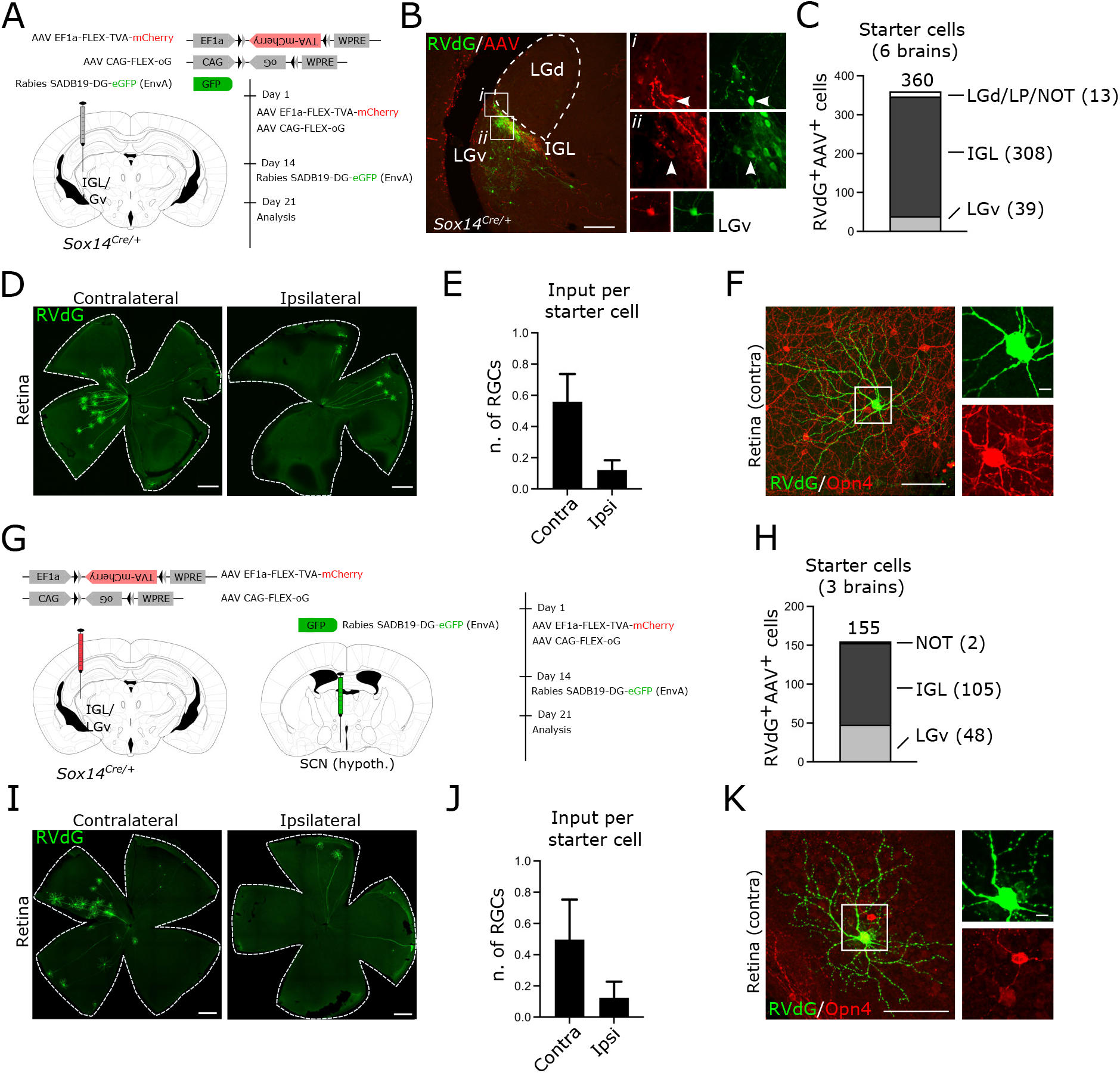
Transsynaptic labelling of retinal input to *Sox14^+^* neurons in the IGL/LGv. ***A***, scheme of rabies tracing of IGL/LGv input in *Sox14^Cre/+^* line, showing the location and timeline of the injections and viral vectors used. ***B***, representative coronal section of virally encoded fluorophores at the injection site, showing RVdG (GFP, green) and helper AAVs (mCherry, red) double-positive neurons within the IGL (white arrowheads) and LGv. Scale bar: 100 μm. ***C***, total count of primary infected cells in the target and off-target regions from serial confocal images of all sections containing thalamic tissue (n = 6 mice). ***D***, representative images of whole mount retinas showing the presence of RVdG-infected RGCs in the retinas ipsilateral and contralateral to the injected IGL/LGv. Scale bars are 500 µm. ***E***, the number of labelled RGCs per starter cell in the ipsi and contralateral retinas (n = 6 retina pairs). ***F***, example image of a retrogradely labelled RGC (green) with weak expression of melanopsin (Opn4, red) in its soma. Note the presence in the same field of view of other RGCs with much higher levels of melanopsin expression. Scale bars: overview image 100 μm, inset 10 μm. ***G***, scheme of rabies tracing of the input to the IGL/LGv subset with hypothalamic projections, showing the location and timeline of viral injections. ***H***, total count of primary infected cells in the target and off-target regions from serial confocal images of all sections containing thalamic tissue (n = 3 mice). ***I***, representative images of whole mount retinas showing the presence of RVdG-infected RGCs in the retinas ipsilateral and contralateral to the injected IGL/LGv upon RVdG injection in the SCN region. Scale bars are 500 µm. ***J***, the number of labelled RGCs per starter cell in the ipsi and contralateral retinas after the injection strategy in **G** (n = 3 retina pairs). ***K***, example image of a retrogradely labelled RGC (green) with weak expression of melanopsin (Opn4, red) in its soma, after RVdG injection in the SCN region. Scale bars: overview image 100 μm, inset 10 μm.

While it is known that M1 and non-M1 classes of melanopsin-type ipRGCs project to the IGL/LGv (Beier et al., 2020; Brown et al., 2010; Ecker et al., 2010; Hattar et al., 2006; Quattrochi et al., 2019; Stabio et al., 2018), it is unclear whether RGC subtype input to the IGL/LGv neurons correlates with the developmental origins of IGL/LGv cells. We therefore set out to investigate the transsynaptic spread of the RVdG to the retina and noted consistent labelling of these cells in the contralateral eye and to a lesser extent in the ipsilateral eye (Fig. 2D). Normalised RGC input was 0.56 ± 0.18 in the contralateral eye and 0.12 ± 0.06 in the ipsilateral eye (Fig. 2E; input per starter cell; mean ± s.e.m, n = 6 mice).

Next, we screened RVdG-infected RGCs for melanopsin expression by immunohistochemical detection of melanopsin on whole mount retinas and noted that none had the high levels of melanopsin expression typically associated with the M1 class; furthermore, dendritic morphologies did not resemble the stereotypical organisation characteristic of the M1 class (Fig. 2F; 261 cells, 12 retinas; n = 6 mice). However, moderate to weak expression was often, but not always, observed in RVdG-labelled RGCs (Fig. 2F).

The absence of any obvious M1 input to the *Sox14^+^* IGL/LGv was unexpected, as *Opn4^taulacZ/+^*;*Sox14^Gfp/+^* mice show that *Sox14^+^* neurons and axons from mostly M1 ipRGCs are clearly present in that same area of the geniculate (Fig. 1M). Furthermore, numerous reports have hypothesised M1 specific innervation of the IGL/LGv (Chen et al., 2011; Ecker et al., 2010; Hattar et al., 2006). Although unlikely, there exist the possibility that our transsynaptic labelling of the RGCs is heavily skewed towards the small number of off target primary infected neurons detected in the LGd, LP and NOT (Fig. 2C), which could explain the absence of M1 input in favour of other melanopsin and non-melanopsin RGC subtypes known to project to thalamic visual areas. We therefore modified the RVdG labelling strategy by delivering the Cre-dependent AAVs in the IGL/LGv, but the RVdG in the ipsilateral SCN (Fig. 1G). While this viral delivery strategy did not result in the exclusive targeting of the SCN, the spread of the injected RVdG viral solution affected only the hypothalamic territory adjacent to the SCN. Hence, by injecting the RVdG in a well-known and distant target of the IGL/LGv we ruled out the possibility of detecting retinal input to the sparse *Sox14^+^* interneurons of thalamocortical nuclei. This modified injection strategy reliably labelled hypothalamus-projecting *Sox14^+^* IGL/LGv, as indicated by the cumulative total number of primary infected neurons from all thalamus-containing sections (3 mice) in the target region (Fig. 2H; IGL: 105, LGv: 48), with residual off target labelling observed in *Sox14^+^* neurons in the NOT region (Fig. 2H; 2 cells). RVdG infected RGCs were detected in all brains analysed (Fig. 2I), with the following normalised distribution: contralateral RGCs 0.50 ± 0.26 and ipsilateral RGCs 0.12 ± 0.1 (Fig. 2J; mean ± s.e.m, n = 3 mice). Inspection of the retinas did not reveal any obvious M1 dendritic morphology, nor was strong melanopsin expression seen in any of the RVdG-infected RGCs (Fig. 2K; 39 cells, n = 3 mice). This observation therefore reinforced our deduction that the thalamic component of the IGL/LGv, including neurons that project to the SCN area of the hypothalamus, lacks an obvious M1 input and suggests that this important source of luminance information may depend on a different and parallel *Sox14^neg^* IGL/LGv circuitry.

To confirm absence of M1 input and to further describe the morphological features of RVdG-labelled RGCs, we quantified radial dendritic morphology and dendritic stratification in the inner plexiform layer (IPL) relative to the ChAT^+^ ON layer and the ChAT^+^ OFF layer of amacrine cell processes (Sumbul et al., 2014) (Fig. 3A). The stratification of the IPL was divided in 10 layers (Rompani et al., 2017; Siegert et al., 2009) so that ChAT^+^ ON and OFF layers match strata 7 and 3, respectively. A custom MATLAB script (kindly provided by Padraic Calpin, UCL) was used to bin the summed arbour density into each of these 10 strata and thresholding applied to colour-code densely populated bins (Fig. 3A; grey). We selected 40 RGCs from animals with AAVs and RVdG injected in the IGL/LGv (6 mice) and 12 RGCs from animals with AAVs injected in the IGL/LGv and RVdG injected in the hypothalamic SCN region (Fig. 3B; n = 3 mice). Criterium for selection of the RGCs was their spatial segregation from other labelled RGCs, so that dendritic morphologies could be reliably reconstructed. We then grouped all reconstructed RGCs into three classes based on whether their dendritic stratification aligned with the ChAT ON lamina (ON), both the ChAT ON and OFF (ON-OFF) or the ChAT OFF lamina (OFF; Fig. 3B,C); hence, the nomenclature adopted is not intended to reflect physiological properties, but dendritic stratification only. Plotting of the dendritic distribution in the IPL for the three groups clearly supports our initial observation of lack of M1 type input, as only 3 cells belonged to the OFF class, none of which was labelled by the hypothalamic injection of the RVdG (Fig. 3C). The ON group contained 21 RGCs, 17 labelled by injecting the RVdG in the IGL/LGv and 4 labelled by injecting it in the SCN hypothalamic region (Fig 3B,C). The ON-OFF group contained 28 RGCs, of which 8 were labelled by the SCN hypothalamic injection of the RVdG (Fig. 3B,C). While, the ON class displayed more homogenous dendritic stratification, the ON-OFF class appears clearly heterogeneous, with bistratified RGCs as well as RGCs showing broad dendritic density spanning the ChAT ON and OFF layers (Fig. 3C). To further characterise the types of RGCs in each of the three groups, we measured the diameter of the dendritic field, the total length of the dendritic tree and the number of dendritic branchpoints (Fig. D,E). We then performed Scholl analysis to measure dendritic complexity at increasing distances from the cell soma (Fig. 3F). We did not measure soma size, because of the heavily saturated GFP signal of the cell somas in our confocal images. RGCs in the ON group have dendritic features consistent with ON stratifying M2, M4 and M5 ipRGCs (Fig. 3D,E,G; range of branchpoints: 18 – 90, range of dendritic tree diameter: 191.2 – 428.5, range of total dendritic length: 2301 – 6821) in line with other reports for these ipRGC subtypes (Berson et al., 2010; Ecker et al., 2010; Estevez et al., 2012; Schmidt and Kofuji, 2011; Stabio et al., 2018). Overall, RGCs in the ON-OFF group displayed broader morphological heterogeneity and significantly larger total dendritic length and branch points compared to the ON group (branch points: p = 0.0005; total dendritic length: p = 0.012; field diameter: p = 0.031; Kruskal-Wallis test). Stratification of the dendritic tree in the IPL discriminates M3 ipRGCs from similarly complex M2 dendrites (Schmidt and Kofuji, 2011). While some of the RGCs in this group display parameters compatible with the M3 type, other RGCs display higher branch points and dendritic length compatible with the recently described M6 ipRGCs (Quattrochi et al., 2019).

**Figure 3.**
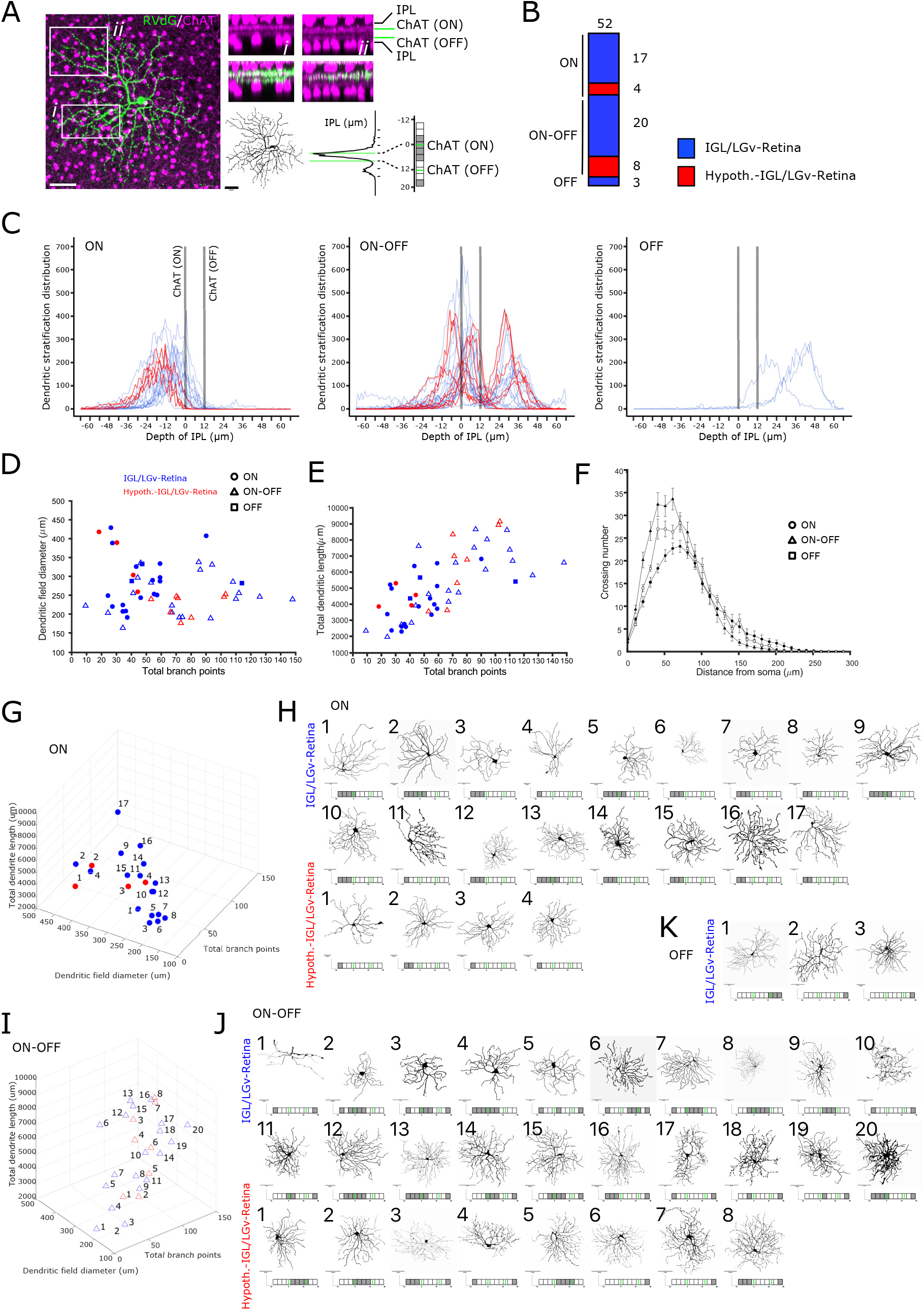
Dendritic morphology of RGCs projecting to *Sox14^+^* neurons in the IGL/LGv. **A**, example images illustrating the strategy for the morphological analysis of RGC dendrites and their stratification in the IPL. The IPL is visualised sandwiched between ChAT^+^ amacrine cell somas (magenta) for two arbitrary regions (*i* and *ii*) of an RVdG infected RGC (green). The dendritic reconstruction of the RGC is presented as black skeleton and the raw density of the GFP signal in the IPL is subdivided along 10 bins using grey colour for thresholding. Green lines represent ChAT^+^ ON and OFF laminar references. Scale bars: 50 μm. ***B***, morphological analysis was conducted for RGCs that did not have overlapping dendrites with other labelled RGCs. Graphical representation of the proportion of ON, ON-OFF and OFF stratifying RGCs, colour coded in blue when labelling occurred after RVdG injection in the IGL/LGv and in red when labelling occurred after RVdG injection in the hypothalamic region of the SCN (cumulative values from 9 mice). ***C***, distribution of dendritic densities for each of the 52 reconstructed RGCs, grouped according to their stratification in the ON and OFF laminas of the IPL and colour coded in blue or red as in **B**. ***D***,***E***, scatter plots displaying the dendritic field diameter and total dendritic length against the total dendritic branch points for all 52 reconstructed RGCs, colour coded in blue or red as in **B**. Filled circles represent the cells in the ON cluster, triangles the cells in the ON-OFF cluster and squares the cells in the OFF cluster. ***F***, Scholl analysis for the 3 clusters of RGCs, without colour coding to indicate location of the RVdG injection. ***G***, 3D rendering of the morphological features in **D**,**E** for the ON RGC cluster, colour coded in blue or red according to the tracing strategy. ***H***, reconstructed dendritic trees and dendritic stratification profiles of retrogradely traced RGCs in the ON cluster, numbered according to their increasing branch points and divided according to the labelling strategy. Scale bars are 50 µm. ***I***, 3D rendering of the morphological features in **D**,**E** for the ON-OFF RGC cluster, colour coded in blue or red according to the tracing strategy. ***J***, reconstructed dendritic trees and dendritic stratification profiles of retrogradely traced RGCs in the ON-OFF cluster, numbered according to their increasing branch points and divided according to the labelling strategy. Scale bars are 50 µm. ***K***, reconstructed dendritic trees and dendritic stratification profiles of retrogradely traced RGCs in the OFF cluster which is only labelled by the RVdG injection in the IGL/LGv. Scale bars are 50 µm.

We cannot exclude that non-ipRGCs are also present in this group, for instance subsets of ON-OFF direction selective (DS) RGCs. The LGv is one of the targets of ON-OFF DS-RGCs (Dhande et al., 2019; Huberman et al., 2009; Rivlin-Etzion et al., 2011) that could potentially mediate some of the non-image forming functions of this RGC class (Rivlin-Etzion et al., 2011).

In summary, reconstruction and quantitative analysis of dendritic morphologies of isolated RGCs retrogradely labelled from *Sox14^+^* IGL/LGv neurons shows that OFF stratifying M1 ipRGCs are not an obvious source of luminance information (Fig. 3C,K), while heterogenous retinal input is mostly from ON (Fig. 3C,G,H) and ON-OFF (Fig. 3C,I,J) stratifying RGCs. Several of the RGCs analysed here have morphological features compatible with non-M1 types of ipRGCs. However, we cannot exclude that other RGCs are also a source of retinal input to the *Sox14^+^* IGL/LGv.

### Brain-wide input to the *Sox14^+^* IGL/LGv is skewed towards visual networks

The IGL/LGv is thought to receive and integrate photic information from the retina with information pertaining to the internal state of an organism, which ascends via the brainstem arousal system. Hence, we systematically analysed the range and proportion of afferents to the *Sox14^+^* neurons of the IGL/LGv by mapping the location of all transsynaptic RVdG-infected cells (GFP^+^) in 4 of the 6 mice used for tracing of the retinal input (Fig. 2A). The vast majority of the afferents arise ipsilaterally in the diencephalic compartments: in the hypothalamus, the zona incerta (ZI) and LGv in the prethalamus, the IGL (including contralateral) and the pHB in the thalamus, the anterior pretectal nucleus (APN) and the nucleus of the optic tract (NOT) in the pretectum (Fig. 4A,B). More than half of the inputs can be grouped under a grossly visual functional classification that includes the retinorecipient subcortical visual shell (Table 4-1; including the superficial layers of the superior colliculus (SCs), the IGL/LGv, the olivary pretectal nucleus (ON) and the NOT), the cortical pyramidal neurons in layer 5 (L5) and 6b (L6b) of the primary visual area (VISp) (Fig. 4A,B) and direct retinal input (Fig. 2D).

**Figure 4.**
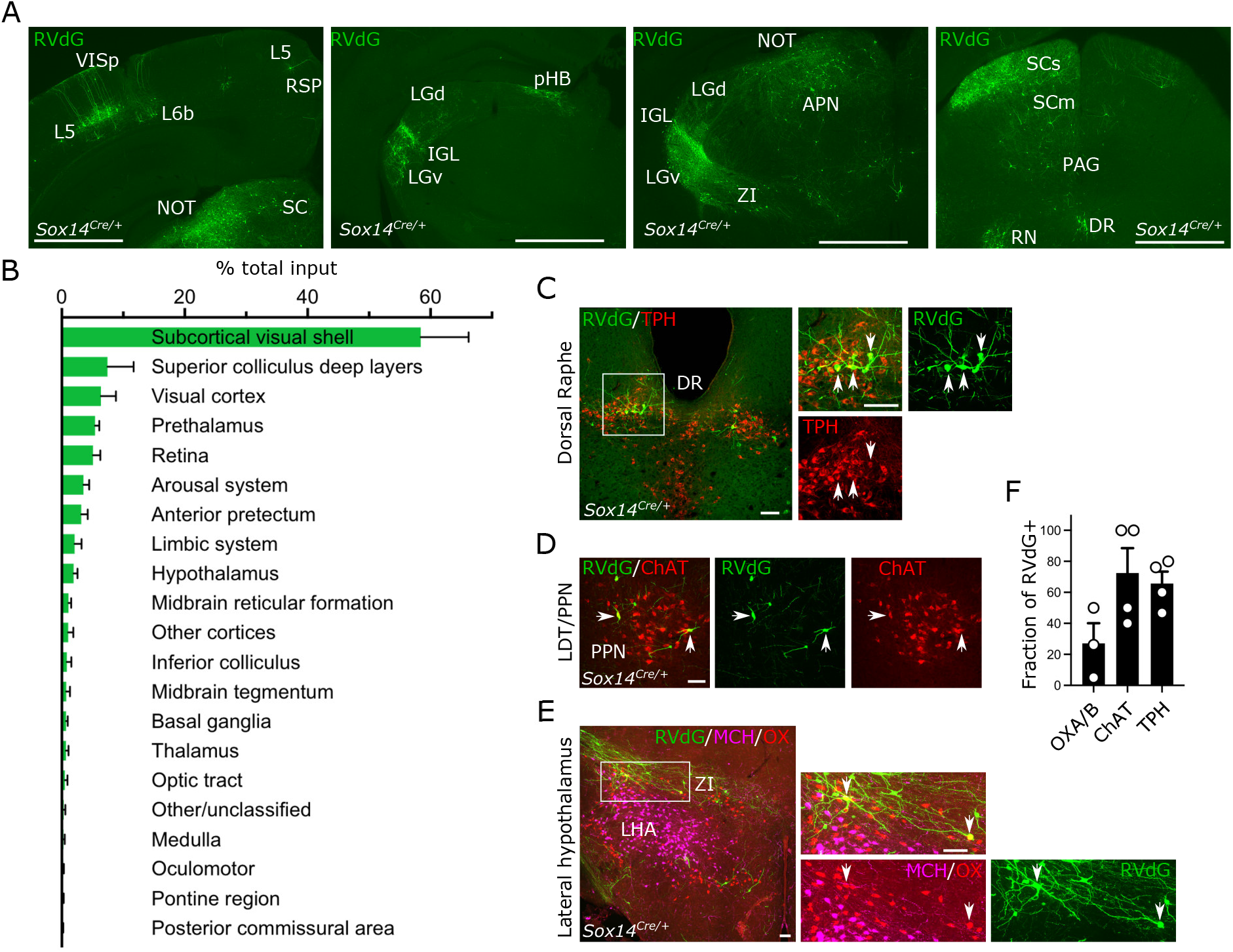
Brain-wide input to the *Sox14^+^* neurons in the IGL/LGv. ***A***, representative coronal sections showing brain-wide distribution of inputs to *Sox14^+^* neurons in IGL/LGv. Most inputs to *Sox14^+^* IGL/LGv can be observed in areas with visual functions (VISp, SCs, NOT, IGL/LGv). Green, GFP from RVdG. Scale bars: ∼1mm. ***B***, quantification of inputs to *Sox14^+^* neurons in IGL/LGv, shown as percentage of total inputs (mean ± s.e.m.; n = 4 mice). See also table 4-1 for anatomical classification based on (Franklin, 2001). ***C***,***D***, representative coronal sections showing inputs from the ascending arousal system. Not all RVdG-infected cells (GFP^+^, green) are co-stained with the anti-tryptophan hydroxylase (TPH, red) antibody in dorsal raphe (DR) or the anti-choline acetyltransferase (ChAT, red) in pedunculopontine nucleus (PPN). Scale bars: 100 mm. ***E***, representative coronal section showing inputs from the lateral hypothalamus. In the lateral hypothalamic area, a small proportion of RVdG infected cells (GFP^+^, green) are co-stained with the anti-orexinA/B antibody (OX, red), but none with the anti-melanin concentrating hormone antibody (MCH, magenta). Scale bars: 100 μm. ***F***, quantification of the fraction of RVdG labelled neurons that expressed the indicated markers (mean ± s.e.m.).

In contrast, input from the ascending arousal system only accounted for less than 5% of the total (Fig. 4B,C,D,E and Table 4-1). Closer inspection revealed this input originates mostly in the serotonergic tryptophan hydroxylase (TPH)^+^ dorsal Raphe (DR; Fig. 4C) and the cholinergic (ChAT^+^) pedunculopontine nucleus (PPN; Fig. 4D). Although input from the locus coeruleus to the IGL has been proposed (Morin, 2013), we could not reliably detect it for the *Sox14^+^* subtype. Furthermore, we noted that not all the RVdG infected neurons in the PPN expressed the cholinergic marker ChAT (Fig. 4F; 72.5 % ± 16.01 %, mean ± s.e.m.). Of the input arising in the DR, 65.7 % ± 7.68 % GFP^+^ cells co-expressed the marker TPH (Fig. 4F; mean ± s.e.m.). In the lateral hypothalamic area (LHA), a small fraction of the retrogradely labelled input was from orexinergic (OX) neurons (Fig. 4F; 27.06 % ± 13.0 %, mean ± s.e.m.) and none was from melanin concentrating hormone (MCH) neurons (Fig. 4E), in agreement to an earlier report in hamsters (Vidal et al., 2005).

In summary, the monosynaptic input to *Sox14^+^* neurons in the IGL/LGv originates overwhelmingly from vision-related structures and to a limited extent from the brainstem’s ascending arousal and neuromodulatory systems.

### The *Sox14^+^* IGL/LGv is required for circadian re-entrainment in presence of weak photic cues

In mammals, alignment of circadian physiology and behaviour with the daily light cycle depends on rod, cones and melanopsin retinal photoreceptors and on specific ipRGC to brain connectivity so that free running circadian rhythms are observed only when all photoreceptors are simultaneously inactivated or ipRGCs selectively ablated (Guler et al., 2008; Hatori et al., 2008; Panda et al., 2003). Melanopsin loss of function mutations alone are not sufficient to cause an overt photoentrainment phenotype, but result in reduced pupillary constriction (Lucas et al., 2003) and reduced behavioural responses to acute light exposure (Altimus et al., 2008; Lupi et al., 2008; Mrosovsky and Hattar, 2003; Panda et al., 2002). The thalamic contribution to circadian photoentrainment is not fully elucidated. Our data showing the notable lack of M1 ipRGC input to the *Sox14^+^* neurons of the IGL/LGv and limited innervation from the ascending arousal system represent unexpected findings that pose the question of the extent to which this developmentally defined subset of the IGL/LGv complex can contribute to circadian entrainment of motor activity rhythms.

We aimed to test the requirement of the IGL/LGv *Sox14^+^* neurons in circadian photoentrainment under normal laboratory lighting conditions (200 lux) and under reduced luminance (10 lux) or photodetection (melanopsin loss of function). To achieve this, we crossed the *Opn4^taulacZ/taulacZ^* and *Opn4^taulacZ/+^*;*Sox14^Cre/+^* mouse lines to generate a *Opn4^taulacZ/+^*;*Sox14^Cre/+^* cohort with functional melanopsin expression and a *Opn4^taulacZ/taulacZ^*;*Sox14^Cre/+^* cohort lacking melanopsin expression. We then induced selective apoptosis of *Sox14^+^* IGL/LGv neurons by injecting bilaterally in the IGL/LGv region an AAV that expresses the diphtheria toxin A subunit in a Cre-dependent manner and the fluorescent reporter mCherry constitutively (AAV2/1 Ef1a-mCherry-DIO-DTA; Fig. 5A). Control animals from both cohorts were injected with a Cre-dependent AAV vector expressing the fluorescent reporter CFP (AAV2/1-Ef1a-DIO-CFP). The extent of ablation was estimated *post hoc* by mapping the spatial extent of fluorophore expression from the AAV vector (Fig. 5B). The successful ablation of *Sox14^+^* neurons in the IGL/LGv was further confirmed by *in situ* hybridisation (ISH) with an RNA probe against the *Npy* mRNA (Fig. 5C; reduction in *Npy^+^* neurons: *Opn4^taulacZ/+^*;*Sox14^Cre/+^* 88 % ± 0.05 %, p < 0.0001; *Opn4^taulacZ/taulacZ^*;*Sox14^Cre/+^* 77 % ± 0.14 %, p = 0.0003; mean ± s.e.m.; unpaired t-test). DTA-ablated and control animals were single-housed in a circadian cabinet, and their spontaneous locomotor activity recorded via passive infrared detectors. Ablation of *Sox14^+^* IGL/LGv neurons did not alter overall rhythmicity of motor behavior nor the circadian period length when animals were housed in constant darkness (tau; Fig. 5D); however a trend towards increased period upon IGL/LGv ablation was noted consistently with previous studies (Pickard, 1994). In both the melanopsin heterozygote (Fig. 5E,G) and knockout background (Fig. 5F,H), ablation of the *Sox14^+^* IGL/LGv neurons did not affect the ability of the mice to photoentrain to standard laboratory light conditions (200 lux;12h light:12h dark; Fig. 5I,N). We then tested the ability of these animals to respond to a 6-hour phase advance in the light cycle, maintaining all other conditions, including luminance, unchanged. Fourteen days after the light phase advance, all groups had entrained to the new light cycle (Fig. 5I,O), indicating that neither the *Sox14^+^* IGL/LGv neurons nor melanopsin expression or both combined are required for circadian resetting of activity rhythms under standard luminance (200 lux). However, ablation of the *Sox14^+^* IGL/LGv appeared to cause a delayed behavioural response to the light phase advance (Fig. 5I), which did not reach statistical significance after correction for multiple comparisons (Fig. 5O; minutes to re-entrainment at day 7: *Opn4^taulacZ/+^Sox14*-control 11.25 ± 11.25 minutes *Opn4^taulacZ/+^Sox14*-ablated: 104.4 ± 17.46 minutes; *Opn4^taulacZ/taulacZ^ Sox14*-control 43.75 ± 25.77 *Opn4^taulacZ/taulacZ^ Sox14*-ablated 119.2 ± 42.43 minutes; mean ± s.e.m.). This observation of delayed entrainment under standard light conditions is consistent with earlier neurotoxin injections in the lateral geniculate of the hamster (Johnson et al., 1989).

**Figure 5.**
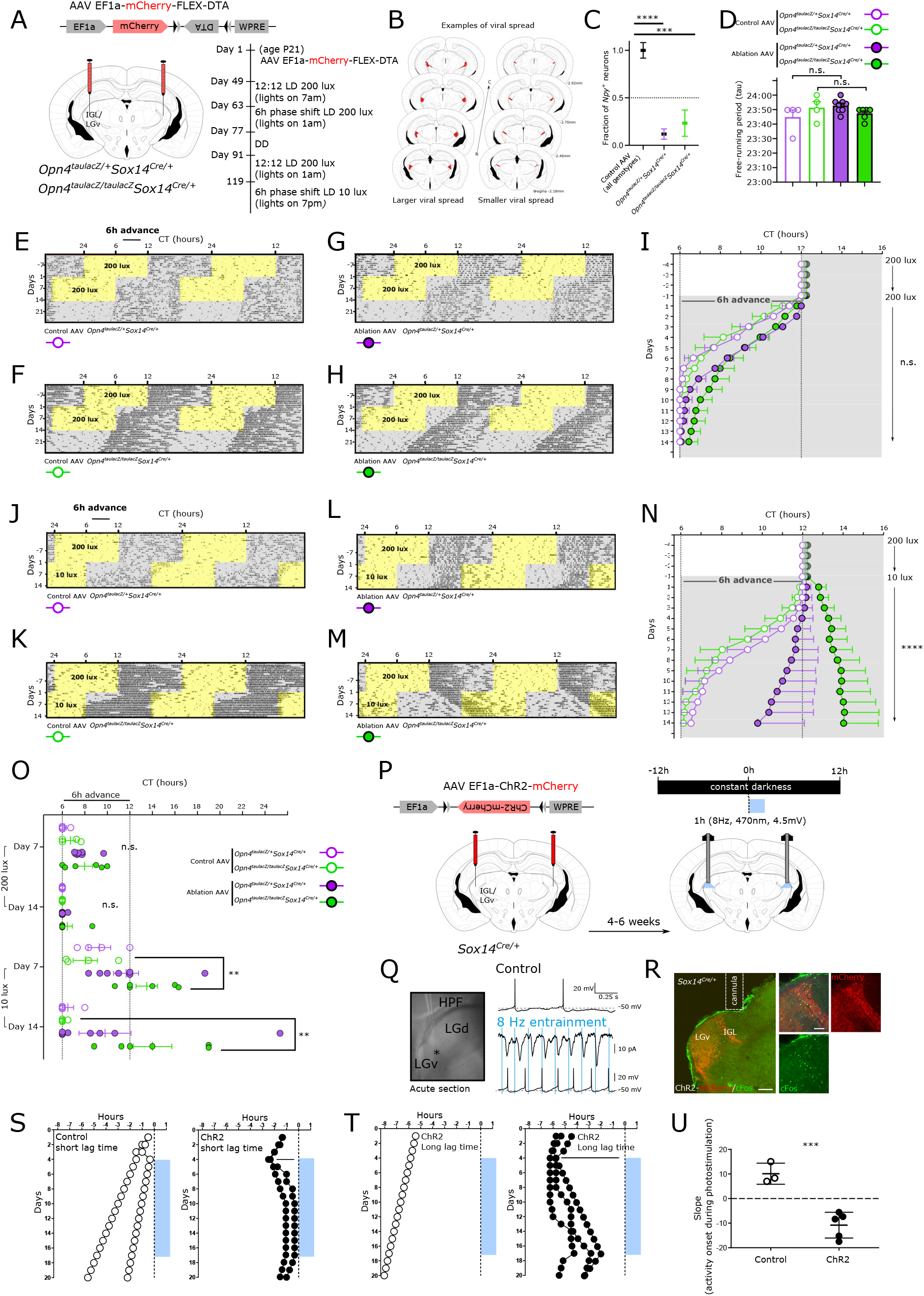
Perturbation of the *Sox14^+^* IGL/LGv neurons leads to aberrant onset of circadian motor activity rhythms. ***A***, scheme for the specific ablation of *Sox14^+^* IGL/LGv neurons in the melanopsin loss of function and control background and timeline of the circadian light paradigms. ***B***, Evaluation of the viral spread upon stereotaxic injection in the IGL/LGv region, in two representative brains depicting the extent of mCherry expression (red), which is not Cre-dependent, in a case of more extensive viral infection (left) and one of less extensive infection (right). ***C***, quantification of the residual fraction of IGL *Npy*^+^ cells, setting the mean value for non-ablated samples at 1. Controls combined: *Opn4^taulacZ/+^Sox14*-control and *Opn4^taulacZ/taulacZ^Sox14*-control *n* = 8 mice, *Opn4^taulacZ/+^Sox14*-ablated *n* = 7 mice, *Opn4^taulacZ/taulacZ^Sox14-*ablated n = 5 mice. One way ANOVA F(32.53). p < 0.0001; mean ± s.e.m. **** < 0.0001; *** = 0.0003; t-test. ***D***, Similar period length under constant dark conditions for all animals in the four groups (n.s.; ANOVA). ***E***,***F***,***G***,***H***, actograms for a representative mouse per group illustrating the activity rhythms under 200 lux, after a 6 hours phase advance and in constant darkness. Period of lights on are in yellow, periods of lights off are in grey. ***I***, onset of circadian motor activity for the four groups, colour coded as in **D**. Periods of lights off are in grey. Days 1 to 14: n.s.; ANOVA. Data plotted as mean ± s.e.m. ***J***,***K***,***L***,***M***, actograms for the same representative mice as in **E**,**F**,**G**,**H** illustrating the activity rhythms under 200 lux, after a 6 hours phase advance with concomitant reduction of ambient luminance to 10 lux. ***N***, onset of circadian motor activity for the four groups. Periods of lights off are in grey. Days 1 to 14: p < 0.0001; Kruskal-Wallis test. *Opn4^taulacZ/taulacZ^Sox14-*ablated significantly different in all pairwise comparisons. Data plotted as mean ± s.e.m.. ***O***, onset of circadian activity rhythms for each animal in the four groups at day 7 and day 14 after the 6 hours phase advance in the light cycle either in standard luminance (200 lux) or dim luminance (10 lux). At 200 lux, a trend towards delayed circadian onset of activity was detected at day 7 however this was not statistically significant (p = 0.041; Kruskal-Wallis test. N.s. after multiple comparison correction). At 10 lux, a high degree of interindividual variability was observed at day 7 and day 14 in the onset of circadian activity rhythms for both the *Opn4^taulacZ/+^Sox14-*ablated and the *Opn4^taulacZ/taulacZ^Sox14-*ablated groups, which included cases of period lengthening and irregular patterns of activity onset (Day 7: F(4.294), *p* = 0.019, ANOVA; Tukey’s multiple comparisons test p = 0.021 for the *Opn4^taulacZ/taulacZ^Sox14-*ablated group. Day 14: p = 0.0082 Kruskal-Wallis test; Dunn’s multiple comparisons test p = 0.021 for the *Opn4^taulacZ/taulacZ^Sox14-*ablated group versus *Opn4^taulacZ/taulacZ^Sox14-*control group and p = 0.028 for the *Opn4^taulacZ/taulacZ^Sox14-*ablated group versus *Opn4^taulacZ/+^Sox14-*control group. Data plotted as mean ± s.e.m.. ***P***, schematic strategy for the expression of ChR2 in *Sox14^+^* IGL/LGv neurons and the circadian optogenetic stimulation (blue line, 1h/day, 470 nm, 8 Hz). ***Q***, a bright-field image of the acute slice preparation indicating the location (asterisk) of whole-cell recordings from mCherry expressing neurons within the IGL/LGv. The top voltage trace was obtained from a single neuron in the IGL/LGv showing spontaneous AP generation in the absence of blue light stimuli. The middle trace shows the ChR2-mediated currents elicited by blue light stimulation and the bottom voltage trace demonstrates the resulting entrainment of APs at 8 Hz optogenetic stimulation. ***R***, representative images from an injected *Sox14^Cre/+^* mouse showing the location of virally infected cells (mCherry, red) in the IGL/LGv region and optogenetically induced c-Fos expression (green), 90 min after the end of the stimulation. Scale bars: 100 µm. ***S***, daily onset of circadian activity in two control subjects (white circle, left graph) and two subjects expressing ChR2 within *Sox14+* IGL/LGv neurons (black circles, right panel), housed under constant dark conditions. Onset of optogenetic stimulation occurred within the first 3 hours of the subjective night. ***T***, daily onset of circadian activity in one control subject (white circle, left graph) and three subjects expressing ChR2 within *Sox14+* IGL/LGv neurons (black circles, right panel), housed under constant dark conditions. Onset of optogenetic stimulation occurred approximately 6 hours into the subjective night. ***U***, plot showing the slope of activity onset over 14 days of optogenetic stimulation in control and ChR2-expressing mice. Positive values indicate negative drifting typical of free running rhythms under constant darkness, whereas negative values reflect a switch to positive drifting. Control group: 10.11 ± 2.48, ChR2-expressing group: −11.38 ± 1.58, p = 0.0002; mean ± s.e.m.; t-test.

We subsequently tested the ability of control and ablated mice to entrain to a further 6-hour phase shift while simultaneously reducing the strength of the light zeitgeber to 10 lux; Fig. 5J,K,L,M,N). Luminance of 10 lux and below mimics the ecologically relevant twilight conditions and suffice to generate SCN activation and behavioural photoentrainment in laboratory mice (Cheng et al., 2004).

Regardless of the status of *Opn4* expression, by day 14 after the dim-light phase advance all but two of the animals that did not experience ablation of *Sox14^+^* IGL/LGv neurons had successfully entrained their activity rhythms to the new dim light cycle (Fig. 5O; minutes to re-entrainment: *Opn4^taulacZ/+^Sox14*-control 30 ± 30 minutes; *Opn4^taulacZ/taulacZ^Sox14*-control 7.5 ± 7.5 minutes, mean ± s.e.m.). Ablation of *Sox14^+^* IGL/LGv impacted the ability of the animals to rapidly entrain to the new dim light cycle (Fig. 5N; p < 0.0001 Kruskal Wallis test).

Ablation of the *Sox14^+^* IGL/LGv had a differential effect on re-entrainment to the dim light cycle depending on the status of *Opn4* expression (Fig. 5J,K,L,M). By day 14 from the dim light phase advance, only 3 out of 8 mice in the *Opn4^taulacZ/+^Sox14*-ablated group had entrained to the new light cycle and one animal displayed a lengthening of the circadian period (positive drifting; Fig. 5O). Furthermore, we observed irregular patters of activity onset (Fig. 5L) during the 14 day-period, which were not present in the same animals exposed to the jet lag paradigm at 200lux. Contrary to the result observed under 200 lux, under dim light the melanopsin loss of function accentuated the entrainment phenotype observed upon ablation of the *Sox14^+^* IGL/LGv (Fig. 5N) so that by day 14, none of the animals in the *Opn4^taulacZ/taulacZ^Sox14*-ablated group had entrained to the dim light cycle and 3 out 6 animals displayed period lengthening (Fig. 5M,N,O; minutes to re-entrainment at day 7: *Opn4^taulacZ/taulacZ^Sox14*-control 122.5 ± 65.11 minutes, *Opn4^taulacZ/taulacZ^Sox14*-ablated: 450.0 ± 56.98 minutes, p = 0.0058, mean ± s.e.m, unpaired t-test; minutes to re-entrainment at day 14: *Opn4^taulacZ/taulacZ^Sox14*-control 7.5 ± 7.5 minutes, *Opn4^taulacZ/taulacZ^Sox14*-ablated 480.8 ± 102.9 minutes, mean ± s.e.m., p = 0.0095, Mann-Whitney test).

Taken together, these data showed that while neither melanopsin expression nor the *Sox14^+^* neurons of the IGL/LGv are required for entrainment and re-entrainment in a jet-lag paradigm, the *Sox14^+^* IGL/LGv neurons contribute to the rapid resetting of circadian activity rhythms. Strikingly, while under luminance levels akin to twilight conditions melanopsin loss of function alone had little effect on the ability of the mice to photoentrain, the combined ablation of the *Sox14^+^* IGL/LGv severely disrupted circadian photoentrainment of activity rhythms.

### Daily optogenetic stimulation of the *Sox14^+^* IGL/LGv neurons entrains motor activity rhythms

Circadian rhythms of nocturnal animals can be entrained by pulses of light given at dusk and dawn, possibly reflective of the light sampling behaviour displayed in their natural ecological niche (DeCoursey, 1986; Edelstein and Amir, 1999; Rosenwasser et al., 1983b; Stephan, 1983b). Consistent with those earlier studies, optogenetics-assisted resetting of circadian oscillatory activity in dark reared mice can be achieved by daily, 1 hour-long, blue light pulses delivered at low frequency (4-8 Hz) on SCN neurons expressing ChR2 (Jones et al., 2015; Mazuski et al., 2018).

To investigate whether experimental activation of *Sox14^+^* IGL/LGv neurons is also sufficient to alter circadian patterns of behaviour, we aimed to replicate this artificial circadian entrainment protocol, stimulating the *Sox14^+^* IGL/LGv neurons instead of the SCN ones. To achieve this, we injected either a Cre-dependent AAV vector expressing the light-gated ion channel Channelrodopsin2 (ChR2; AAV2/5 Ef1a-DIO-hChR2(H134R)-mCherry) or a control AAV expressing the cyan fluorescent protein (AAV2/1-Ef1a-DIO-CFP; Fig. 5P) bilaterally in the IGL/LGv region of *Sox14^Cre/+^*mice. We then tested the impact of forced ChR2-mediated activation of the *Sox14^+^* IGL/LGv neurons for 1 hour at daily intervals on the onset of circadian locomotor activity in animals housed under constant darkness (Fig. 5P).

Pulses of blue light (470nm) were delivered bilaterally directly above the IGL/LGv at a frequency of 8 Hz (Fig. 5P), which falls within the physiological frequency range of IGL neurons (Chrobok et al., 2021; Chrobok et al., 2018; Lewandowski and Blasiak, 2004) and of the SCN neurons during the light phase of the day (Jones et al., 2015; Sakai, 2014). Furthermore, optogenetic 10 Hz stimulation of RGC axon terminals is sufficient to activate IGL neurons (Shi et al., 2020).

Ex vivo patch clamp recording from ChR2-expressing IGL neurons confirmed reliable light-induced responses to 8 Hz blue light entrainment (Fig. 5Q). In vivo, this optogenetic protocol led to expression of the immediate early gene c-Fos in the IGL/LGv (Fig. 5R), which is a reliable marker of neuronal activation (Dragunow and Faull, 1989; Peters et al., 1996).

As expected, housing under constant dark conditions efficiently induced free running rhythms in control and experimental mice (Fig. 5S,T,U; white and black circles, respectively). However, while daily optogenetic light stimulation in control animals had no significant impact on the onset of circadian motor activity (Fig. 5S,T; white circles), it affected it profoundly in experimental animals that expressed ChR2 (Fig. 5S,T; black circles). The impact of daily stimulation of the *Sox14^+^* IGL/LGv is reflected in the drastic change in the slope of the circadian onset of motor activity between the two groups (Fig. 5U; p = 0.0002; t-test).

The gradual shift of the activity onset, which moved progressively towards the time of stimulation and in some but not all cases, locked onto it, is strikingly similar to the results obtained when stimulation was applied directly onto the SCN (Jones et al., 2015; Mazuski et al., 2018). Acute light pulses during the active phase in nocturnal animals caused negative masking of motor activity (Morin and Studholme, 2014a; Mrosovsky et al., 2001; Redlin, 2001), a response that has recently been suggested to depend on IGL neurons (Shi et al., 2020). In our experimental conditions, a reduction in spontaneous motor activity was detected during the first episode of optogenetic stimulation of the *Sox14^+^* IGL/LGv neurons (motor activity change during optogenetic stimulation compared to hour following it ChR2 group: 38.51 % ± 19.72 %; CFP group: 391.4 % ± 123.0 %, mean ± s.e.m., p = 0.0093; t-test); however the decrease in motor activity was negligeable over seven consecutive days of optical stimulation (ChR2 group: 141.0 % ± 13.76 %; CFP group: 229.8 % ± 37.48 %, mean ± s.e.m., p = 0.036; t-test).

The effect of the optogenetic stimulation of the *Sox14^+^* IGL/LGv neurons on the phase of circadian motor activity was observed regardless of the lag time between the endogenous onset of locomotor activity and the time of the optogenetic stimulation (Fig. 5S, short lag time and Fig. 5T, long lag time). This optogenetically-induced effect on the onset of circadian motor activity is rapidly reversed upon termination of the photostimulation (Fig. 5S,T). Hence, in absence of a strong zeitgeber such as circadian light, daily optogenetic stimulation of the *Sox14^+^* neurons of the IGL/LGv is sufficient to reorganise circadian locomotor activity.

### The *Sox14^+^* IGL/LGv neurons are required for rapid change in vigilance states at circadian light transitions

Circadian transitions between light and dark regulate neuronal network dynamics that contribute to shaping the sleep-wake cycle. We hypothesised that the broad innervation of the *Sox14^+^* neurons in the IGL/LGv by visual networks may reflect an underappreciated role in regulating rapid changes in vigilance in response to circadian light transitions. Such brain network changes may not be readily detected by monitoring gross circadian locomotor activity but can be more reliably measured as changes in the spectral power of a cortical electroencephalogram (EEG).

To monitor the impact of ablating the *Sox14^+^* IGL/LGv neurons on the vigilance states of the brain at each light transition under standard circadian conditions, we injected male *Sox14^Cre/+^* mice with either AAV Ef1a-mCherry-DIO-DTA (ablated group) or AAV Ef1a-DIO-CFP (control group) into the IGL/LGv region, replicating the genetic strategy previously described. Control and ablated mice were then implanted with screw type skull electrodes for EEG (Fig. 6-1) and stainless-steel wire type electrodes inserted into the trapezius muscle of the neck for electromyogram (EMG). During the EEG/EMG recording, animals could move freely in their cage. Both ablated and control animals displayed characteristic cortical spectrograms, EEG/EMG traces and hypnograms with detectable transitions between periods of high magnitude delta frequency oscillations and reduced mobility indicative of non-rapid eye movement (NREM) sleep (Fig. 6A,B), and high magnitude theta frequency oscillations occurring either without associated increase in the amplitude of the EMG signal, as is typical of rapid eye movement (REM) sleep, or with associated EMG activity, as is typical of the wake state (Wake; Fig. 6A,B).

**Figure 6.**
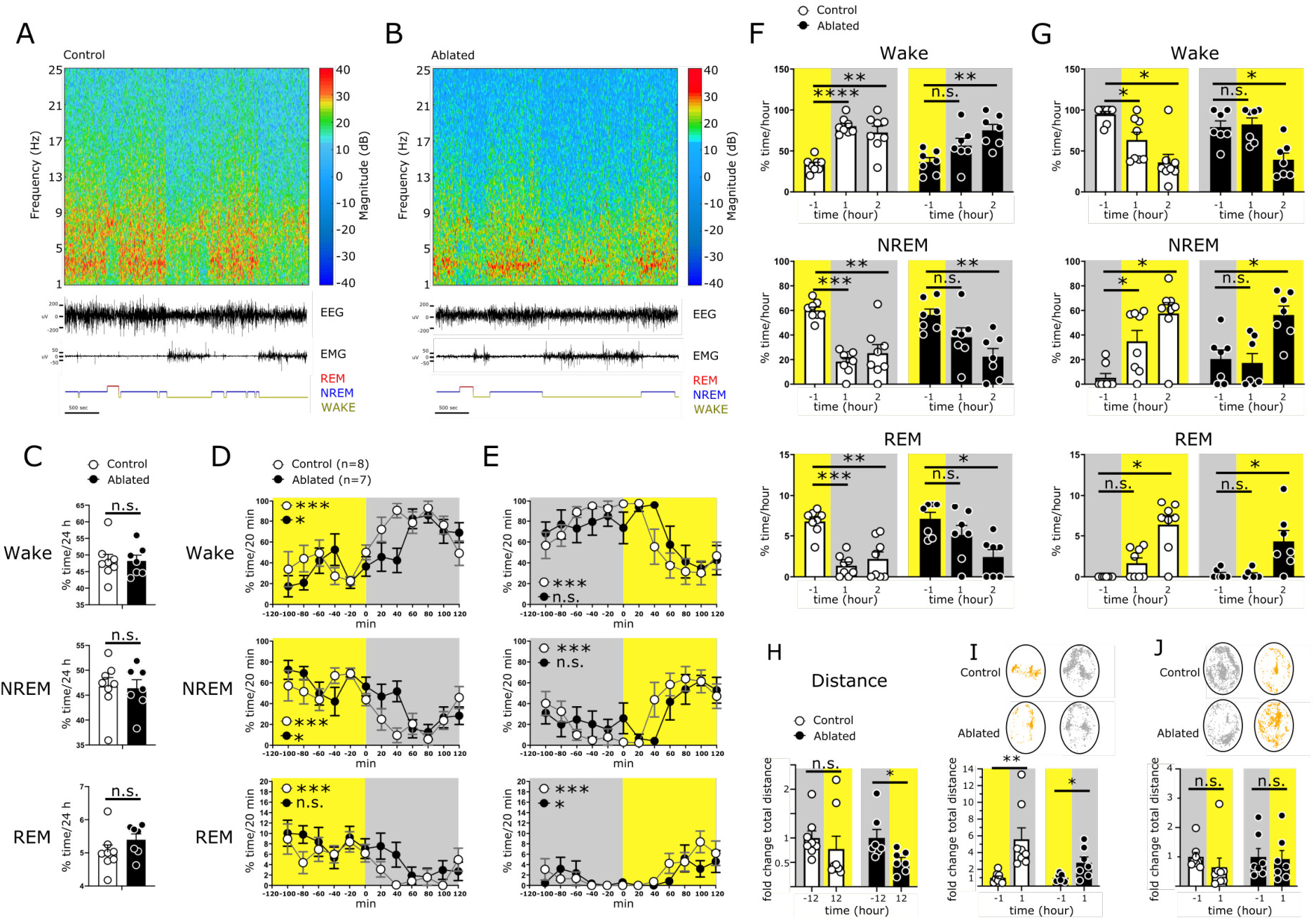
Ablation of *Sox14^+^* IGL/LGv neurons causes delayed vigilance state transitions at circadian light changes. ***A***,***B***, spectrograms, EEG/EMG traces, and hypnograms over a one-hour period from a representative control mouse and an ablated mouse. ***C***, over a 24-hour period control and *Sox14^+^* IGL/LGv-ablated mice spend a similar percentage of time in Wake (48.24 % ± 1.96 % vs 48.22 % ± 1.71 %, p = 0.99), NREM (46.73 % ± 1.80 % vs 46.38 % ± 1.71 %, p = 0.89) and REM (5.03 % ± 0.20 % vs 5.39 % ± 0.17 %, p = 0.2). Values are mean ± s.e.m., t-test. ***D***,***E***, distribution of Wake, NREM and REM in the two hours preceding and following each light transition. Light and dark hours are shaded in yellow and grey, respectively (see Table 6-1 for comprehensive statistics). ***F***, pairwise comparison in the content of Wake, NREM and REM between the hour preceding and each of the two hours following the light to dark transition. Control group: Wake_-1_: 33.17 % ± 2.98 %, Wake_1_: 80.26 % ± 3.49 %, p < 0.0001, Wake_2_: 72.61 % ± 7.76 %, p = 0.001; NREM_-1_: 60.08 % ± 2.65 %, NREM_1_: 18.39 % ± 3.29 %, p = 0.0001, NREM_2_: 25.17 % ± 7.04 %, p = 0.0015; REM_-1_: 6.75 % ± 0.52 %, REM_1_: 1.35 % ± 0.45 % p = 0.0004, REM_2_: 2.18% ± 0.85 % p = 0.007. Ablated group: Wake_-1_ 36.57 % ± 5.34 %, Wake_1_: 56.67 % ± 3.49 %, p = 0.093, Wake_2_: 75.23 % ± 7.24 %, p = 0.008; NREM_-1_ 56.35 % ± 4.73 %, NREM_1_: 38.19 % ± 7.68 %, p = 0.093, NREM_2_: 22.36 % ± 6.64 %, p = 0.0081; REM_-1_: 7.08 % ± 0.78 %, REM_1_: 5.10 % ± 1.16 %, p = 0.296, REM_2_: 2.41 % ± 0.90 %, p = 0.031. Values are mean ± s.e.m., t-test except Control REM_-1_ versus REM_2_, Ablated REM_-1_ versus REM_1_ and REM_2_ Wilcoxon test. ***G***, pairwise comparison in the content of Wake, NREM and REM between the hour preceding and each of the two hours following the dark to light transition. Control group: Wake_-1_: 94.77 % ± 3.48 %, Wake_1_: 63.47 % ± 9.42 %, p = 0.015, Wake_2_: 36.11 % ± 9.72 %, p = 0.015; NREM_-1_: 5.22 % ± 3.48 %, NREM_1_: 34.87 % ± 8.82 % p = 0.015, NREM_2_: 57.51 % ± 8.86 % p = 0.015; REM_-1_: 0.00 % ± 0.00 %, REM_1_: 1.66 % ± 0.64 %, p = 0.125, REM_2_: 6.37 % ± 1.09 %, p = 0.015. Ablated group: Wake_-1_ 79.18 % ± 7.51 %, Wake_1_: 82.47 % ± 7.80 %, p = 0.812, Wake_2_: 39.37 % ± 8.09 %, p = 0.017; NREM_-1_ 20.50 % ± 7.34 %, NREM_1_: 17.25 % ± 7.65 %, p = 0.848, NREM_2_: 56.31 % ± 7.22 %, p = 0.021; REM_-1_: 0.31 % ± 0.21 %, REM_1_: 0.27 % ± 0.20 %, p > 0.999, REM_2_: 4.31 % ± 1.34 %, p = 0.0313. Values are mean ± s.e.m., Wilcoxon test except Ablated Wake_-1_ versus Wake_2_ and NREM_-1_ versus NREM_2_ t-test. ***H***, fold change in distance travelled in the light phase (12 hours). Total distance in the dark phase (12 hours) was set at 1. Control group: 1.0 ± 0.14 (dark) versus 0.77 ± 0.26 (light); Ablated group: 1.0 ± 0.17 (dark) versus 0.52 ± 0.82 (light). ***I***, Example trajectories for one control and one ablated mouse in the hour preceding and following the light change. Oval: ROI used for automated tracking; yellow: light, grey: dark. Histograms report the fold change in distance travelled in the hour preceding and following the light change. For the light to dark transition distance in the light was set at 1: Control group 1.0 ± 0.21 (light) 5.59 ± 1.36 (dark), p = 0.0078; Ablated group 1.0 ± 0.17 (light) 2.80 ± 0.68 (dark), p = 0.044. ***J***, For the dark to light transition distance in the dark was set at 1: Control group 1.0 ± 0.15 vs 0.64 ± 0.31, p = 0.148; Ablated group 1.0 ± 0.28 vs 0.92 ± 0.29, p = 0.468. Values are mean ± s.e.m., Wilcoxon test except for Ablated light to dark transition t-test. A summary of statistical tests is available in Table 6-1.

Consistent with the pattern of circadian locomotion observed in the cohort of *Sox14^+^* IGL/LGv-ablated mice tested for circadian light entrainment, the cumulative fraction of a 24-hour period spent in NREM, REM and Wake states did not differ between ablated and control groups (Fig. 6C and Table 6-1; NREM: p = 0.89, REM: p = 0.2, Wake: p = 0.99), indicating that the *Sox14^+^* neurons of the IGL/LGv are not required for sleep and wake overall. However, plotting the content of NREM, REM and Wake for each hour of the circadian cycle revealed a discrepancy between ablated and control groups specifically in the hour following each light transition. We further increased the temporal resolution by binning NREM, REM and Wake episodes in 20 minutes intervals for the 2 hours preceding and following each light transition. This analysis revealed that, while both control and ablated animals increased the time spent in Wake in the first 2 hours after lights off (Fig. 6D and Table 6-1 for full statistical data; Wake Control: p < 0.0001; Wake Ablated: p = 0.015) and conversely, decreased it at lights on (Fig. 6E and Table 6-1; Wake Control: p < 0.0001, Wake Ablated: p = 0.051), the ablated group remained for a protracted period in a state similar to the one preceding each light transition, responding with a delayed kinetic to the circadian light change (Fig. 6D,E).

To quantitatively assess the impact of circadian light transitions on Wake, NREM and REM states upon ablation of the *Sox14^+^* IGL/LGv, we plotted the cumulative time spent in each of the three vigilance states during the hour preceding the light change (−1) and the first (1) and the second hour (2) after the light change for the control and ablated groups (Fig. 6F). In the control group, pairwise comparisons between the hour preceding and each of the two hours following the light transition showed a strong and statistically significant change in each of the three vigilance states already in the first hour following the circadian light transition (Fig. 6F and Table 6-1; Wake_-1_ versus Wake_1_: p < 0.0001; NREM_-1_ versus NREM_1_: p = 0.0001; REM_-1_ versus REM_1_: p = 0.0004). In contrast, change in all three vigilance states in the ablated group does not reach statistical significance during the first hour after the light transition (Fig. 6F and Table 6-1, Wake_-1_ versus Wake_1_: p = 0.093; NREM_-1_ versus NREM_1_: p = 0.093; REM_-1_ versus REM_1_: p = 0.296). However, in the ablated group, change in Wake and NREM reached statistical significance in the second hour after the light transition (Fig. 6F and Table 6-1; Wake_-1_ versus Wake_2_: p = 0.0080; NREM_-1_ versus NREM_2_: p = 0.0081), substantiating the interpretation that the *Sox14^+^* IGL/LGv neurons are required for the rapid change in cortical network activity caused by circadian light transitions, but not for overall regulation of sleep and wake over a 24-hour period.

We then investigated whether a similar requirement for the *Sox14^+^* IGL/LGv neurons also exists at the dark to light transition. In the control group, the time spent in Wake, NREM and REM over the 3-hour period changed significantly (Table 6-1 for full statistical data).

In keeping with the observed rapid changes in vigilance states at light to dark transition (Fig. 4F), rapid changes in Wake and NREM were also detected already in the first hour after the transition to lights-on (Fig. 6G and Table 6-1; Wake_-1_ versus Wake_1_: p = 0.015; NREM_-1_ versus NREM_1_: p = 0.015; REM_-1_ versus REM_1_: p = 0.125). However, in the ablated group, significant changes in the three vigilance states only became apparent in the second hour after the light transition (Fig. 6G and Table 6-1; Wake_-1_ versus Wake_2_: p = 0.017; NREM_-1_ versus NREM_2_: p = 0.021; REM_-1_ versus REM_2_: p = 0.031). Hence, *Sox14^+^* IGL/LGv neurons are required at both circadian light transitions to elicit rapid transitions between vigilance states.

While delayed responses to the circadian light transitions caused by the ablation of *Sox14^+^* IGL/LGv neurons could readily be detected in the EEG/EMG data, we noted that they correlated less clearly with overt motor behaviour. Change in the distance travelled before and after the circadian light transitions, assessed by automated video tracking (Pinnacle Technologies, Inc.) of animals undergoing EEG/EMG recordings, did not reveal similarly striking changes within and between groups (Fig. 6H,I,J and Table 6-1).

We therefore focused our analysis on cortical network activity rather than behavioural output and extended the analysis of the EEG data to quantify oscillatory activity in the delta (0.5-4Hz), theta (6-9Hz) and alpha (8-12Hz) frequency ranges. High power delta oscillations are typically observed during NREM, while theta oscillations are associated with REM sleep as well as exploratory behaviour and alpha oscillations are enriched during quiet wake.

Power spectral densities displayed dynamic changes at either circadian light transition in both the control (Fig. 7A,G) and the ablated group (Fig. 7D,J). In the control group, time-frequency analysis of EEG data showed the presence of delta waves dominating the hour preceding the light to dark transition, with the appearance of theta and low alpha waves anticipating the light change and a sharp decrease in delta following the transition (Fig. 7A). The clear shift from delta to theta waves in the control group was supported by a theta/delta ratio (T/D) that was significantly increased after the circadian light change (Fig. 7B and Table 7-1; (T/D)_-1_ versus (T/D)_1_ p = 0.0016) and the clear frequency shift from high delta and low theta state in the hour preceding the light transition (Fig. 7C, yellow line) to high theta and low delta state in the hour following the light transition (Fig. 7C, grey line).

**Figure 7:**
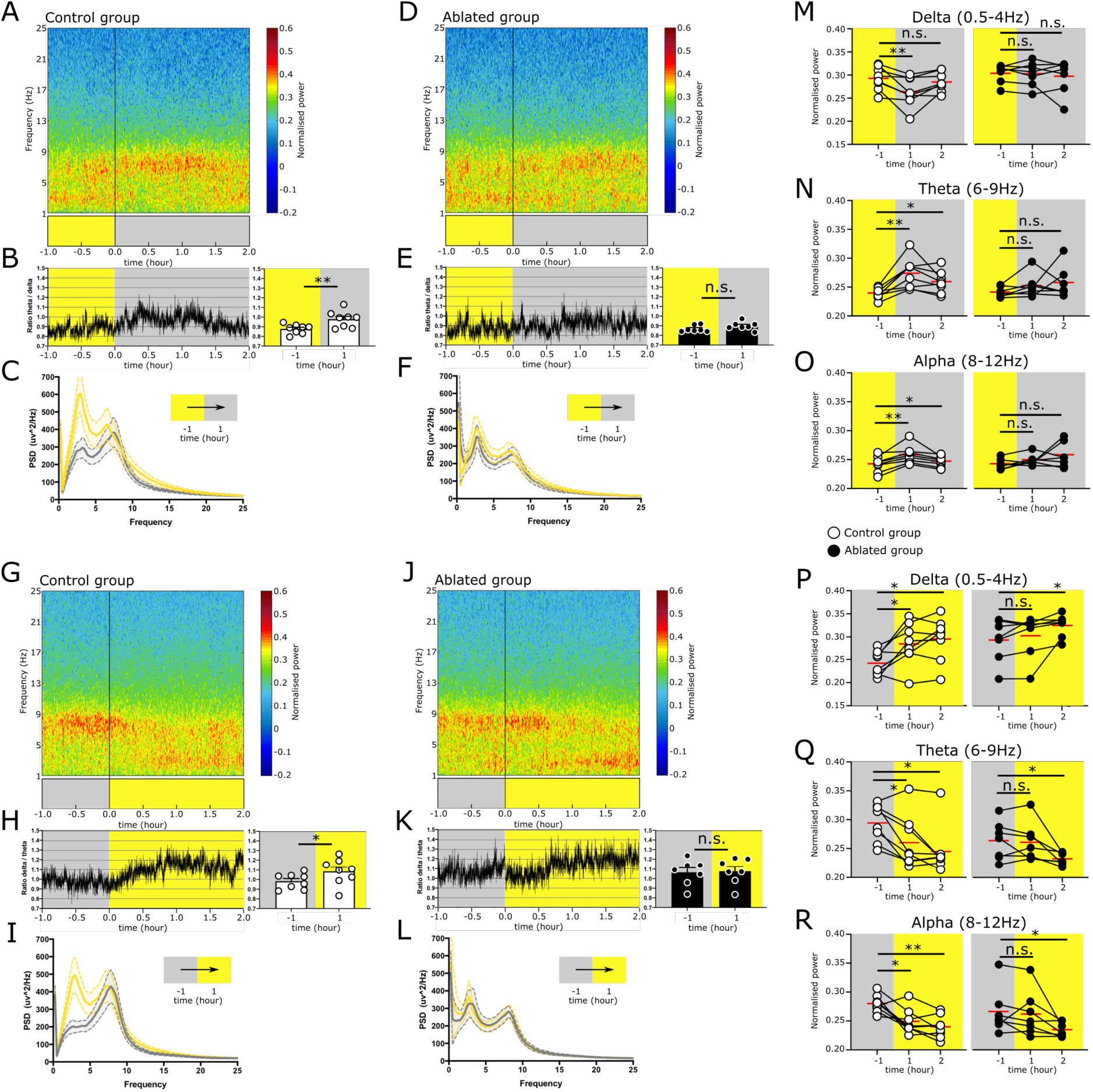
The *Sox14^+^* IGL/LGv neurons are required for the rapid onset of cortical network activity associated with circadian light transitions. ***A***, averaged cortical spectrogram for the Control group displaying a rapid increase in the theta power and decrease in delta power at the transition from lights on to lights off. ***B***, the change in theta and delta is displayed as a group average theta/delta ratio over time and as cumulative for the hour preceding and the hour following the light change. Control group (T/D)_-1_: 0.87 ± 0.01 versus (T/D)_1_: 0.97 ± 0.03, p = 0.001. Values are mean ± s.e.m., t-test. ***C***, the power spectrum of the Control group in the hour preceding the transition from light to dark is overlayed to the power spectrum profile for the hour following the light transition (bold line: mean; dotted lines: ± s.e.m.; yellow: light, grey: dark). ***D***, averaged cortical spectrogram for the Ablated group displaying retention of high delta power following the lights on to lights off transition. ***E***, the change in theta and delta is displayed as a group average theta/delta ratio over time and as cumulative for the hour preceding and the hour following the light change. Ablated group (T/D)_-1_: 0.86 ± 0.01 versus (T/D)_1_: 0.89 ± 0.01, p = 0.078. Values are mean ± s.e.m., Wilcoxon test. ***F***, the power spectrum of the Ablated group in the hour preceding the transition from light to dark is overlayed to the power spectrum profile for the hour following the light transition (bold line: mean; dotted lines: ± s.e.m.; yellow: light, grey: dark). ***G***, averaged power spectrum for the Control group in the hour preceding and the two hours following the lights off to lights on transition. ***H***, the change in the delta/theta ratio is plotted over time and as group mean for the hour preceding and following the light transition. Control group (D/T)_-1_: 0.98 ± 0.02 versus (D/T)_1_: 1.08 ± 0.04, p = 0.016. Values are mean ± s.e.m, t-test. ***I***, the power spectrum of the Control group in the hour preceding the transition from dark to light overlayed to the power spectrum profile for the hour following the light transition (bold line: mean; dotted lines: ± s.e.m.; yellow: light, grey: dark). ***J***,***L***, the same analysis as in G-H is applied to the Ablated group. Note that the Ablated group does not display a significative change in the delta/theta ratio. Ablated group (D/T)_-1_: 1.06 ± 0.05 versus (D/T)_1_: 1.07 ± 0.05, p = 0.623. Values are mean ± s.e.m., *t*-test. ***M***, pairwise comparison for the delta power for the hour preceding and each of the two hours following the transition from lights on to lights off in the Control group (white circles) and Ablated group (filled black circles). Control group Delta_-1_: 0.29 ± 0.009, Delta_1_: 0.26 ± 0.01 p = 0.009, Delta_2_: 0.28 ± 0.007 p = 0.323. Values are mean ± s.e.m., t-test. Ablated group Delta_-1_: 0.30 ± 0.007, Delta_1_: 0.30 ± 0.009 p = 0.734, Delta_2_: 0.29 ± 0.013 p = 0.937. Values are mean ± s.e.m., Wilcoxon test. ***N***, pairwise comparison for the theta power for the hour preceding and each of the two hours following the transition from lights on to lights off in the Control group (white circles) and Ablated group (filled black circles). Control group Theta_-1_: 0.23 ± 0.003, Theta_1_: 0.27± 0.009, p = 0.002, Theta_2_: 0.25± 0.006, p = 0.105. Values are mean ± s.e.m., t-test. Ablated group Theta_-1_: 0.24 ± 0.003, Theta_1_: 0.25± 0.007 p = 0.109, Theta_2_: 0.25± 0.01, p = 0.109. Values are mean ± s.e.m., Wilcoxon test. ***O***, pairwise comparison for the alpha power for the hour preceding and each of the two hours following the transition from lights on to lights off in the Control group (white circles) and Ablated group (filled black circles). Control group Alpha_-1_: 0.24 ± 0.004, Alpha_1_: 0.25 ± 0.005, p = 0.003, Alpha_2_: 0.24 ± 0.003, p = 0.045. Values are mean ± s.e.m., t-test. Ablated group Alpha_-1_: 0.24 ± 0.003, Alpha_1_: 0.24 ± 0.003, p = 0.109, Alpha_2_: 0.25 ± 0.008, p = 0.097. Values are mean ± s.e.m., Wilcoxon test. ***P***,***R***, similar pairwise analysis as in M-O but for the transition from lights off to lights on. Control group Delta_-1_: 0.24 ± 0.009, Delta_1_: 0.28± 0.016, p = 0.009, Delta_2_: 0.29 ± 0.016, p = 0.019. Ablated group Delta_-1_: 0.29 ± 0.017, Delta_1_: 0.30 ± 0.017, p = 0.296, Delta_2_: 0.32 ± 0.009, p = 0.036. Values are mean ± s.e.m., t-test except for the Ablated group Delta_2_ comparison which used a Wilcoxon test. Control group Theta_-1_: 0.29 ± 0.011, Theta_1_: 0.26 ± 0.016, p = 0.023, Theta_2_: 0.24 ± 0.014, p = 0.023. Values are mean ± s.e.m., Wilcoxon test. Ablated group Theta_-1_: 0.26 ± 0.012, Theta_1_: 0.26 ± 0.012, p = 0.706, Theta_2_: 0.23 ± 0.003, p = 0.038. Values are mean ± s.e.m., t-test. Control group Alpha_-1_: 0.27 ± 0.005, Alpha_1_: 0.24 ± 0.007, p = 0.012, Alpha_2_: 0.23 ± 0.006, p = 0.005. Values are mean ± s.e.m, t-test. Ablated group Alpha_-1_: 0.26 ± 0.014, Alpha_1_: 0.26± 0.014, p = 0.375, Alpha_2_: 0.23 ± 0.004, p = 0.046. Values are mean ± s.e.m., Wilcoxon test. A summary of statistical tests is available in Table 7-1.

In stark contrast, the spectrogram of EEG data from the *Sox14^+^* IGL/LGv ablated mice showed a mixture of delta, theta and low-alpha waves in the hour before the light transition. Delta waves continued to be present for the first hour after the light transition, eventually fading in the second hour, when theta and low-alpha oscillations increased (Fig. 7D). Consequently, there was no significant change in T/D ratio (Fig. 7E; (T/D)_-1_ versus (T/D)_1_: p = 0.078) and power spectral densities showed similar spectral patterns before (Fig. 5F, yellow line) and after the light transition (Fig. 7F, grey line). These observations are consistent with delayed cortical network dynamics at the lights-on to lights-off transition in mice with *Sox14^+^* IGL/LGv ablation.

We then performed the same analysis for the lights-off to lights-on transition of the circadian day, which is normally accompanied by sleep onset. As expected, spectrograms from the control mice showed the presence of theta and low-alpha waves in the hour preceding the light transition, which shifted to high amplitude delta and reduced theta and low-alpha waves already within the first hour following the light transition (Fig. 7G). This clear shift from theta to delta waves in the control group was reflected in the increased delta/theta ratio (D/T) (Fig. 7H and Table 7-1; (D/T)_-1_ versus (D/T)_1_: p = 0.0168).

Similarly, power spectral density showed a clear frequency shift from high amplitude theta and low delta in the hour preceding the light transition (Fig. 7I, grey line) to high delta and low theta within the first hour following the light transition (Fig. 7I, yellow line).

The ablated mice differed from the control group as theta and low alpha waves persisted for approximately 40 minutes following the light transition, after which they faded and were replaced by delta oscillations (Fig. 7J). Consequently, there was no significant change in D/T ratio in the hour preceding and following the light transition (Fig. 7K; (D/T)_-1_ versus (D/T)_1_: p = 0.623) and power spectral densities retained similar distribution before (Fig. 7L, grey line) and after the light transition (Fig. 7L, yellow line).

To assess quantitatively the change in power for the delta, theta and alpha ranges, we calculated the cumulative power for the hour preceding and for the first and second hour following each circadian light transition, starting first from lights-on to lights-off. Across the three time points, animals in the control group showed significant change for all three frequency bands (Fig. 7M,N,O; Delta p = 0.0064, Theta p = 0.0012, Alpha p = 0.0031). Delta power decreased significantly already in the first hour after the light transition (Fig. 7M and Table 7-1; Delta_-1_ versus Delta_1_ p = 0.0099). Concomitantly, the theta power increased significantly (Fig. 7N and Table 7-1; Theta_-1_ versus Theta_1_: p = 0.0027) as did alpha power (Fig. 7O and Table 7-1; Alpha_-1_ versus Alpha_1_: p = 0.0039). In contrast, the group of animals with ablation of the *Sox14^+^* neurons in the IGL/LGv did not display significant overall change in the power of delta, theta and alpha across the three timepoints (Fig. 7M,N,O and Table 7-1 for comprehensive statistical data).

We then performed the quantitative analysis on the combined hourly variation in delta, theta and alpha for the second circadian light transition of the day, corresponding to lights off to lights on. As expected, significant difference was detected in the control group (Delta: p = 0.011, Theta: p = 0.030, Alpha: p = 0.0029). Pair-wise comparisons between the hour preceding and each of the two hours following the circadian light change confirmed a significant increase in delta power taking place already in the first hour (Fig. 7P and Table 7-1; Delta_-1_ versus Delta_1_: p = 0.019), accompanied by a significant decrease in theta power (Fig. 7Q and Table 7-1; Theta_-1_ versus Theta_1_: *P* = 0.023) and alpha power (Fig. 7R and Table 7-1; Alpha_-1_ versus Alpha_1_: p = 0.0124).

In contrast, in the *Sox14^+^* IGL/LGv ablated group, delta and alpha power did not change significantly across the three time points (Fig. 7P,R and Table 7-1; Delta: p = 0.051, Alpha: p = 0.111). Power in the Theta frequency across all time points showed a significant difference (Table 7-1; Theta: p = 0.028), which was due to a decrease in power in the second hour after the transition (Fig. 7Q and Table 7-1; Theta_-1_ versus Theta_1_: p = 0.706, Theta_-1_ versus Theta_2_: p = 0.038).

In summary, power spectral analysis of cortical EEG reveals a previously undescribed role for the thalamic neurons in the IGL/LGv in enabling rapid changes in cortical network activity at both circadian light changes.

### *Sox14^+^* neurons in the IGL/LGv and pHB are related

In mice, thalamic GABA projection neurons are also present in the pHB (An et al., 2020). We and others have previously described *Sox14^+^* GABAergic precursors that during embryonic development migrate tangentially from the presumptive IGL/LGv to reach the location of the presumptive pHB (Fig. 1A), indicating a developmental lineage relationship between GABA projection neurons in these two thalamic structures (Delogu et al., 2012; Vue et al., 2007). To test whether *Sox14^+^* neurons in the IGL/LGv and the pHB retain shared patterns of connectivity, we adapted the transsynaptic RVdG strategy previously used to map synaptic input to the *Sox14^+^* neurons of the IGL/LGv to the *Sox14^+^* cells in the pHB (Fig. 8A,B). Anatomical mapping of the input to the *Sox14^+^* neurons of the pHB revealed an overall striking similarity with the input to the *Sox14^+^* neurons of the IGL/LGv with a large fraction arising in the subcortical visual shell (Fig. 8F and Table 4-1). In contrast to the *Sox14^+^* IGL/LGv neurons, the *Sox14^+^* neurons of the pHB did not receive detectable retinal input, nor input from the primary cortical visual area. However, they displayed a sizeable input (∼20%) from the IGL/LGv (Fig. 8C,D,E) and from subcortical and cortical limbic structures (Fig. 8C,F and Table 4-1).

**Figure 8.**
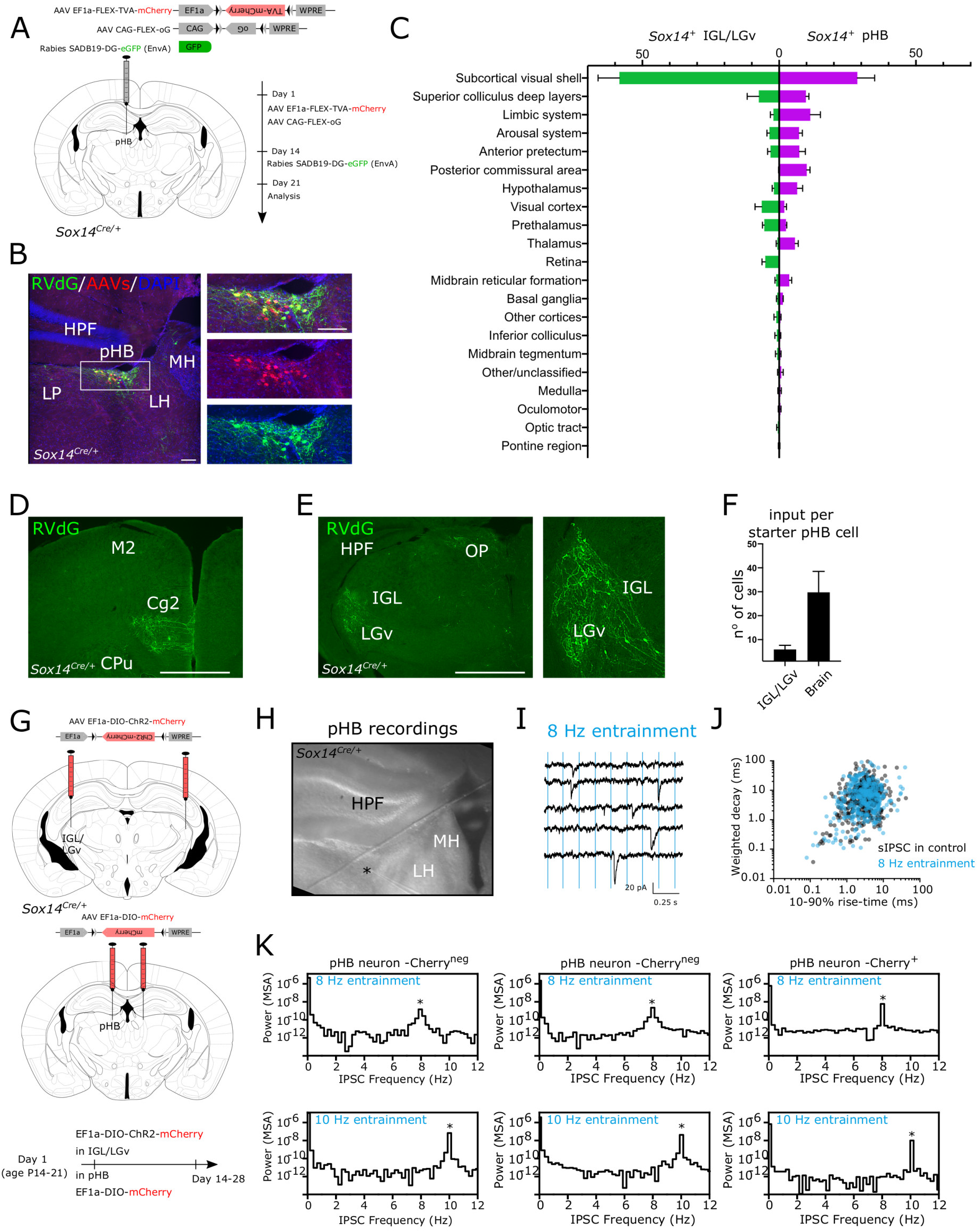
Developmentally related *Sox14^+^* neurons in the IGL/LGv and pHB retain a large proportion of shared input. ***A***, scheme of rabies tracing in *Sox14^Cre/+^* line, showing the location of the injection site (pHb) and the viruses used. ***B***, representative coronal section of the injection site, showing starter cells (GFP^+^RFP^+^ double-positive neurons) within the pHB. Green, GFP from RVdG; red, mCherry from AAVs. HPF: hippocampal formation, LP: lateral posterior, LH: lateral habenula, MH: medial habenula. Scale bar 100 μm. ***C***, quantification of inputs to *Sox14^+^* neurons of pHB (right, n = 3 mice) from anatomically defined regions, compared to input to the *Sox14^+^* neurons in the IGL/LGv (left, n = 4 mice) from the same anatomical regions. Data are shown as percent of total number of cells (mean ± s.e.m.). ***D***, representative coronal sections of the brain-wide distribution of inputs to *Sox14^+^* neurons of pHB. M2: secondary motor cortex, Cg2: cingulate cortex area 2, CPu: caudate putamen. Scale bar 100 μm. ***E***, representative coronal section at thalamic level showing the presynaptic neurons in the IGL/LGv innervating the *Sox14^+^* neurons in the pHB. HPF: hippocampal formation, OP: olivary pretectal nucleus. Scale bar 100 μm. ***F***, the normalised IGL/LGv input (5.93 ± 1.68) is a sizable fraction of all brain wide afferents (29.77 ± 8.75) to the pHB. Inputs are normalised for the number of starter cells in the pHB. mean ± s.e.m.; n = 3 mice. ***G***, strategy to introduce ChR2 in *Sox14^+^* IGL/LGv neurons and mCherry in *Sox14^+^* pHB neurons. ***H***, example of the acute brain preparation illustrating the location of a patched cell. ***I***, postsynaptic currents in pHB neurons from an acute preparation upon 8Hz light stimulation of ChR2-expressing axonal terminals. ***J***, peristimulus time histogram analysis of sIPSCs. ***K***, examples of entrained activity in pHB neurons with or without mCherry expression upon laser stimulation at 8 Hz and 10 Hz.

These data complement our observation of monosynaptic input from the area of the pHB to the *Sox14^+^* neurons of the IGL/LGv (Fig. 4B) and suggest that the thalamic *Sox14^+^* neurons of the IGL/LGv and the pHB retain a large proportion of shared input, but with specific differences such as retinal and prefrontal limbic inputs. Following AAV delivery of ChR2 (EF1a-DIO-ChR2-mCherry) to neurons of the IGL/LGv in the *Sox14^Cre/+^* mouse (Fig. 8G) we attempted to record from mCherry expressing neurons within the pHB (Fig. 8H) that had been injected with a second AAV (EF1a-DIO-mCherry). The very low number of viable mCherry-expressing neurons within the pHB hindered functional identification of monosynaptic connectivity between IGL/LGv and pHB. However, peristimulus time histogram (PSTH) analysis of sIPSC timing (Fig. 8I,J) in mCherry^+^ and mCherry^neg^ neurons of the pHB (n = 3 mice) supports the presence of entrained activity resulting from optogenetic stimulation of the Sox14^+^ IGL/LGv input (Fig. 8K).

Intrigued by the possibility of an IGL/LGv input to the pHB, we aimed to further characterise the *Sox14^+^* neurons in the pHB by identifying their axonal projections and compare these with those of other recently described pHB cell types such as the ventral pHB neurons that projects to the ventromedial prefrontal cortex (vmPFC) (An et al., 2020; Fernandez et al., 2018) and the GABAergic neurons of the dorsal pHB that project to the nucleus accumbens (An et al., 2020). To achieve this, we injected in the pHB of *Sox14^Cre/+^* mice either a cre-dependent AAV expressing a cell membrane-bound form of GFP (AAV2/1 Ef1a-DIO-mGFP; Fig. 9A) or a presynaptic-localised Synaptophysin-GFP fusion (AAV2/1 phSyn1(S)-FLEX-tdTomato-SypEGFP; Fig. 9D). The AAV-labelled *Sox14^+^* pHB neurons (Fig. 9B) occupy an area with dense axonal projections from the *Sox14^+^* IGL/LGv (Fig. 9B’). The axonal projections of the *Sox14^+^* pHB extend along two main directions: rostro-ventral, along the lateral thalamus to terminate in the IGL/LGv and medial, penetrating the habenular complex (Fig. 9B,C) and terminating predominantly in the medial habenula (MH; Fig. 9C,E,F,G). The presynaptic terminals of *Sox14^+^* pHB neurons are enriched in a Calb2^+^ Chat^neg^ domain in the dorsal portion of the MH (Fig. 9E,F) and, in keeping with the shared developmental origin with the thalamic component of the IGL/LGv, contained the GABA transporter vGat (Fig. 9G). Hence, our structural and functional connectivity data support a model whereby tangential migration of *Sox14^+^* precursors from the developing IGL/LGv to the pHB enables formation of a thalamic network of GABAergic neurons which may represent a previously undescribed substrate for thalamic limbic functions.

**Figure 9.**
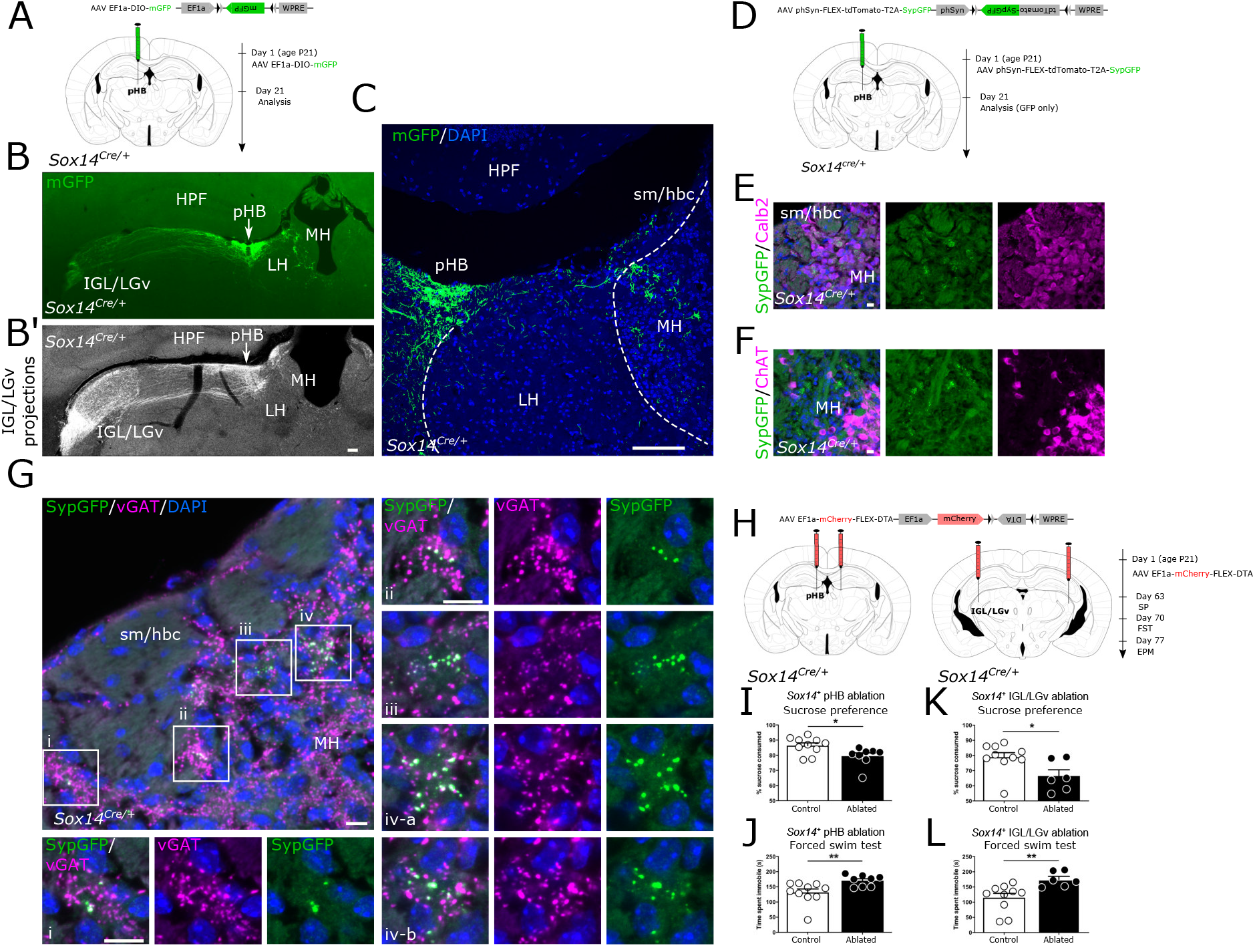
The *Sox14*^+^ IGL/LGv and pHb neurons are part of a thalamic network for mood regulation. ***A***, scheme of injection of Cre-dependent AAV expressing mGFP in pHB of *Sox14^Cre/+^* mice. ***B***,***B’***, representative coronal sections showing the reciprocal projections between the IGL/LGv and the pHB. In B’, for comparison the projections of the *Sox14^+^* IGL/LGv. Green, mGFP. Scale bar 100 μm. ***C***, representative coronal section showing the injection site and the second route of projections from the pHB, terminating in the MH. Green, mGFP. Sm: stria medullaris, hbc: habenular commissure. Scale bar 100 μm. ***D***, scheme of injection of Cre-dependent AAV expressing SypGFP in pHB of *Sox14^Cre/+^* mice. ***E***,***F***,***G***, representative coronal sections showing the synaptic terminals (GFP^+^) of *Sox14^+^* pHB neurons in the MH. Most sypGFP positive terminals are within the Calb2^+^ portion of the MH (**E**) and anti-vGAT (**G**) antibodies in MH, but not with the anti-ChAT antibody (**F**). Green, SypGFP; magenta, Calb2 (**E**), ChAT (**F**), vGAT (**G**). Scale bar 10 μm. ***H***, scheme for specific ablation of *Sox14^+^* pHB and IGL/LGv neurons using an AAV-encoded DTA-expression cassette and behavioural tests’ timeline (see also Figure 9-1). ***I***,***K***, *sucrose preference test*. Both *Sox14^+^* pHB-and *Sox14^+^* IGL/LGv-ablated mice consumed less sucrose solution compared to control subjects. PHB control group: 86.38 % ± 1.77 %, pHB ablated group: 79.40 % ± 2.19 %, p = 0.021. IGL/LGv control group: 78.84 % ± 2.99 %, IGL/LGv ablated group: 66.26 % ± 4.05 %, p = 0.042. Data are expressed as mean ± s.e.m., U-test. ***J***,***L***, *Forced swim test*. Both pHB-and IGL/LGv-ablated mice spent more time immobile compared to control subjects. PHB control group: 131.6 ± 11.39 s, pHB ablated group: 170.1 ± 6.56 s, p = 0.006. IGL/LGv control group: 114.8 ± 14.22 s, IGL/LGv ablated group: 175.1 ± 10.07 s, p = 0.002. Data are expressed as mean ± s.e.m., U-test.

### The *Sox14^+^* neurons of the IGL/LGv and pHB participate in mood homeostasis

Recent reports have shown how both the IGL/LGv and pHB participate in mood regulation (An et al., 2020; Fernandez et al., 2018; Huang et al., 2019). The discovery of a projection from *Sox14^+^* pHB neurons to the dorsal MH is intriguing, because the MH participates in the modulation of ventral midbrain dopaminergic and serotonergic neurons via the fasciculus retroflexus projection to inhibitory interneurons in the rostral part of the interpeduncular nucleus (Lima et al., 2017).

We hypothesised that the *Sox14^+^* neurons in the IGL/LGv and pHB may be part of a previously undescribed circuitry for mood regulation. As proof of principle of a shared participation of the two groups of thalamic *Sox14^+^* neurons in reward pathways, we performed a targeted ablation of the *Sox14^+^* neurons in the pHB by stereotaxic viral delivery of DTA or control CFP in *Sox14^Cre/+^* male mice, as previously described (Fig. 9H and Fig. 9-1). We then assessed behavioural changes in adult *Sox14^+^* pHB ablated mice by evaluating anhedonia, using the sucrose preference test (SP) and behavioural despair, using the forced swim test (FST), while maintaining animals in standard single housing conditions.

Mice with ablation of the *Sox14^+^* neurons of the pHB showed a decreased sucrose preference (Fig. 9I; p = 0.021) and an increased time spent immobile in the FST (Fig 9J; p = 0.006) compared to control subjects. This finding is consistent with the *Sox14^+^* pHB participating in maintenance of mood tone, potentially acting through their projection to the MH. We then replicated the ablation and behavioural testing strategy for the *Sox14^+^* neurons of the IGL/LGv to assess whether this would result in a similar change in the SP and the FST, in animals kept under standard single housing conditions. Also in this case, IGL/LGv-ablated mice showed decreased sucrose preference compared to control animals (Fig. 9K; p = 0.042) and spent significantly more time immobile in the FST than control animals (Fig. 9L; p = 0.002). In contrast, anxiety-related behaviour measured as the time spent in the open arms of the elevated-plus maze (EPM) test displayed no significant difference in both pHB-and IGL/LGv-ablated cohorts (pHB control: 33.12 ± 9.07 s, pHB ablation: 27.35 ± 3.7 s, p = 0.598 and IGL/LGv control: 17.85 ± 3.53 s, IGL/LGv ablation: 30.7 ± 7.14 s, p = 0.09, mean ± s.e.m., t-test). Our observations add to recent work implicating the IGL/LGv (Huang et al., 2019) and the pHB (An et al., 2020; Fernandez et al., 2018) in mood regulation by suggesting that developmentally related *Sox14^+^* neurons in the IGL/LGv and the pHB regions participate in mood homeostasis.

## Discussion

While the *Sox14^+^* neurons constitute a thalamic cell class prominently found in the IGL, they also contribute to the prethalamic LGv nucleus and thalamic pHB nucleus. Several fate mapping experiments have demonstrated that LGv neurons arise from prethalamic progenitors (Delaunay et al., 2009; Golding et al., 2014; Inamura et al., 2011; Puelles et al., 2020; Suzuki-Hirano et al., 2011; Vue et al., 2007). Here, we have shown that prethalamic progenitors make a significative contribution to the IGL, a structure of the thalamus proper. The specific circuit organisation of the developmentally defined lineages that make up the IGL/LGv complex is largely unknown. Our monosynaptic restricted retrograde tracing places the *Sox14^+^* IGL/LGv firmly within the visual system, with input from cortical and subcortical visual structures. Innervation of the *Sox14^+^* neurons from the cholinergic, monoaminergic and orexinergic neurons appeared more limited, contrasting with previous classic tract tracing experiments of the anatomically defined IGL/LGv complex (Morin, 2013). It would be intriguing to test if input to prethalamic lineages of the IGL/LGv displayed a complementary shift towards the ascending arousal system.

Previous anterograde tracing of retinal input from *Opn4^+^* RGCs has provided clear evidence of enriched ipRGC innervation of the IGL/LGv from multiple ipRGC types (Chen et al., 2011) as well as conventional RGC types (Beier et al., 2020). Tracing with an M1-enriched reporter construct identified the IGL, which receives afferents from both the *Brn3b^+^* and the *Brn3b^neg^* ipRGCs (Chen et al., 2011), as a major target of the M1 ipRGC subtype (Ecker et al., 2010; Hattar et al., 2006). Surprisingly, our monosynaptic retrograde tracing from *Sox14^+^* IGL/LGv did not label any M1 ipRGCs, but highlighted input from ipRGC types that express low levels of *Opn4* and may be more reliant on classic photoreceptors for luminance detection than the M1 type (Sonoda and Schmidt, 2016). As the vast majority of ipRGCs projecting to the SCN are the *Brn3b^neg^* M1 type (Baver et al., 2008; Chen et al., 2011), which also participate in innervation of the IGL/LGv (Chen et al., 2011), it is likely that some neurons in the IGL/LGv receive a copy of the SCN’s retinal input (Pickard, 1985), however, here we show that the *Sox14^+^* neurons of the IGL/LGv complex do not participate in such M1-driven circuitry. Intact retinohypothalamic connectivity and preservation of a hypothetical M1 ipRGC driven IGL/LGv tract to the SCN would ensure circadian photoentrainment upon ablation of the *Sox14^+^* IGL/LGv, unless the strength of the photic cue is reduced, as observed in our experimental conditions. Our findings are consistent with the hypothesis put forward by Pickard that neurons of the geniculohypothalamic tract may convey information on illumination intensity to the SCN (Pickard et al., 1987). The connectional bias of the *Sox14^+^* IGL/LGv towards subcortical and cortical visual networks bears implications for the view that integration of photic and non-photic circadian cues takes place in IGL/LGv neurons to provide a unified output to the SCN. Our data could also be compatible with the presence of segregated pathways to the SCN via different subtypes of IGL/LGv neurons, with the prethalamic component of the IGL/LGv a plausible candidate. Under a circadian dim light cycle, the observed high inter-individual variability of the activity onset in *Sox14^+^* IGL/LGv-ablated mice with or without melanopsin expression, may be explained by the unmasking of one or more *Sox14^neg^* circuitries for internal state-dependent modulation of the circadian clock.

The use of dim light of amplitudes comparable to dusk and dawn carries ethological value, as nocturnal animals in their natural environment are more likely to sample light from a dark burrow and such transient exposure suffice in providing photoentrainment. Indeed, studies on nocturnal flying squirrels (*Glaucomys Volans)* and other rodents that made use of a den cage, have shown that in nocturnal animals the light sampling behaviour appears to be under circadian control, at dusk and dawn (Pratt and Goldman, 1986; Twente Jr., 1955). It is likely that in their ecological niche nocturnal animals are exposed to just few minutes of light each day (DeCoursey, 1986). Accordingly, regular pulses of light at dusk and dawn are sufficient to photoentrain behavioural rhythms in nocturnal animals (DeCoursey, 1972; Rosenwasser et al., 1983a; Stephan, 1983a). Here we have shown, using an optogenetics strategy, that repeated circadian activation of *Sox14^+^* neurons in the IGL/LGv interrupts the spontaneous drifting of endogenous activity rhythms in mice kept under constant darkness and results in entrainment of the activity onset, in striking similarity to the effect caused by an analogue optogenetics stimulation of the SCN (Jones et al., 2015; Mazuski et al., 2018).

In nocturnal rodents, acute light exposure causes negative masking of motor activity, a process that involves rapid NREM induction (Lupi et al., 2008; Morin and Studholme, 2009, 2014a; Mrosovsky et al., 2001). The subcortical networks involved in this response to acute light exposure are not fully mapped but are thought to involve the pretectum and superior colliculus (Miller et al., 1998; Zhang et al., 2019). More recently, a genetic strategy to ablate GABAergic neurons in the IGL region resulted in reduced NREM sleep upon acute light presentation in mice (Shi et al., 2019), implicating the IGL/LGv complex as an important node in the phenomenon of photosomnolence. It remained unclear whether the IGL/LGv is also required for vigilance state changes at recurrent and predictable circadian light transitions. IpRGCs, with their connectivity to the SCN, but also directly to other brain regions, play an important role in the control of sleep and arousal (Altimus et al., 2008; Lupi et al., 2008; Muindi et al., 2013; Pilorz et al., 2016; Rupp et al., 2019; Tsai et al., 2009). Circadian light transitions initiate a cascade of events that likely involves multiple brain networks and results in the stabilisation of a new vigilance state. Here, we show that the thalamic *Sox14^+^* neurons of the IGL/LGv ensure rapid transition to NREM-associated Delta frequency cortical oscillations at lights-on and, conversely, the rapid establishment of a wake cortical profile at lights-off. Our data substantiate and expand earlier speculative hypotheses implicating the IGL in sleep regulation (Horowitz et al., 2004; Morin, 2013, 2015; Morin and Blanchard, 2005). However, the precise downstream events that are elicited by *Sox14^+^* IGL/LGv neurons at circadian light transitions remain to be fully elucidated.

Defective encoding of circadian light changes emerges as one of the themes from the cell ablation approach presented here. This is consistent with the failure to photoentrain circadian activity rhythms described in the constitutive *Sox14* knockout mice (Delogu et al., 2012); however, while ablation of *Sox14^+^* IGL/LGv neurons in the mature brain caused delayed transitions in vigilance states at circadian light changes, defective photoentrainment was only detectable under reduced strength of the lighting cues. It is likely that *Sox14* loss of function during brain development across neurons of the subcortical visual shell causes widespread changes in network connectivity and neuronal function that result in more overt changes in arousal and circadian behaviours.

In earlier work, we and others demonstrated that during embryonic development, the pTH-R contributes to the formation of a thalamic structure at the edge of the epithalamus (Delogu et al., 2012; Vue et al., 2007) via tangential migration of *Sox14^+^* neurons from the prospective IGL/LGv (Delogu et al., 2012). Hence, the recently described pHB (An et al., 2020; Fernandez et al., 2018) contains, among other cells, neurons that are developmentally related to the IGL/LGv. The *Sox14^+^* neurons of the pHB receive a distinctive input from the prefrontal cortex, but lack the prefrontal projections and the nucleus accumbens projections described for other neuronal types of the pHB (An et al., 2020; Fernandez et al., 2018); furthermore, we did not identify an obvious direct retinal input to the *Sox14^+^* pHB neurons, but described instead an input from the IGL/LGv complex, which are themselves innervated by ipRGCs. Concomitantly, we have shown that the *Sox14^+^* neurons of the pHB present a potential novel pathway for modulation of mood, via a GABAergic input to the dorsal MH.

While the IGL/LGv was recently shown to mediate light-dependent mood control via a direct projection to the LH neurons (Huang et al., 2019), we noted that the dorsal projection of the *Sox14^+^* IGL/LGv appears to be mostly directed to the pHB region, consistent with the existence of a novel pathway from the retina to the dorsal MH via the IGL/LGv and pHB. Indeed, ablation of the *Sox14^+^* neurons in either the IGL/LGv or pHB impaired mood-related behaviour to a similar extent.

By taking a developmentally informed approach, we have identified specific contributions of thalamic neurons of the IGL/LGv and pHB that expand our understanding of thalamic function in sensory perception, the control of vigilance states and the modulation of mood. While this approach reduced the developmental complexity of the anatomically defined IGL/LGv to reveal some of its unique connectional and functional properties, it does not resolve the further differentiation of the *Sox14^+^* developmental lineage into subsets of molecularly defined mature neurons. Future investigations that exploit single cell genomics and connectomics will reveal the finer grain of parallel and integrated pathways occurring on thalamic projection GABAergic neurons.

## Materials and Methods

### Animals

All mice were kept in the animal facilities of King’s College London. The *Sox14^Cre/+^* (Jager et al., 2016) (MGI ID: MGI:5909921) and *Sox14^GFP/+^* mouse lines (Crone et al., 2008) (MGI ID: 3836003) were maintained in the C57Bl/6 background. The *Opn4^taulacZ^* mouse line (Hattar et al., 2002) (MGI ID: MGI:2449781) was a mixed B6/129 background and was crossed to the *Sox14^Cre/+^* or *Sox14^GFP/+^* mouse lines. The *Dlx5/6^Cre^* (Monory et al., 2006) (JAX Stock No: 008199; MGI ID:3758328) and the *Rosa26^lsl-nuclearGFP^* (Mo et al., 2015) (JAX Stock No: 021039; MGI ID: 5443817) were maintained in the C57Bl/6 background. Experimental procedures were approved by the Ethical Committee for Animal Use of King’s College London and were covered by a Project Licence under the UK Home Office Animals (Scientific Procedures) Act 1986. Mice were kept under normal housing conditions (7am lights on and 7pm lights off, with food and water *ad libitum*), unless otherwise stated for behavioural experiments. All behavioural experiments were performed on adult (> 6 weeks of age) male mice. Tract tracing experiments were performed using animals of both sexes.

### Generation of EnvA-pseudotyped, glycoprotein-deleted rabies virus

The EnvA-pseudotyped, glycoprotein-deleted rabies virus (ΔG-SADB19-eGFP, EnvA; abbreviated RVdG) was produced in house following an established protocol (Osakada and Callaway, 2013). The ΔG-SADB19-eGFP (generous gift from Prof Roska, FMI, Basel, Switzerland) was amplified on BHK-SadGly-GFP cell culture (generous gift from Prof Tripodi, LMB, Cambridge, UK) at 37°C – 3.5% CO_2_ to slow cell cycle. The BHK-EnvA cell line (generous gift from Prof Tripodi) was used for pseudotyping the virus with EnvA envelope protein (grown at 37°C −5% CO_2_). Filtered (Steriflip, 0.22um, Millipore) supernatant was stored at 4°C, concentrated by ultracentrifugation, resuspended in sterile PBS and stored at −80°C in single use aliquots.

### Brain stereotaxic surgeries

Briefly, mice were placed in a digital stereotaxic frame (World Precision Instruments) under 2.5% isoflurane anaesthesia. For viral delivery, the skull was exposed by a midline scalp incision, and the stereotaxic frame was aligned using Bregma and Lambda as visual landmarks. A 33-gauge steel needle was placed above the skull and a hole drilled through the skull bone to expose the brain. Virus solutions (100-250 nl) were injected using a borosilicate glass needle (0.58 OD/ID mm, World Precision Instruments) connected to an air injector (Narishige) or a a Nanoject III (Drummond Scientific) injection system. The following general coordinates were used for IGL/LGv with litter-specific finer adjustments: from Bregma AP = −2.25mm; L = ±(2.20-2.40) mm; DV = −2.85 mm; for the pHB from Breagma AP = −1.6mm; L = ±0.35 mm; DV = −(2.42-2.27) mm); for SCN from Bregma AP = −0.5 mm; L = 0.15 mm; DV = - (5.0-5.25) mm. The glass needle was left in place for an additional 8 min before being slowly removed. Following injection, skin was closed using biocompatible tissue glue (VetBond).

For optogenetics experiments, two cannulas (200 µm core diameter; Doric Lenses) holding optical fibres were inserted and extended to the ventral edges of the dorsal part of the LGN and further fixed to the skull with dental cement. Mice were allowed to recover in a heating chamber and returned to their home cage after waking up. All mice received a subcutaneous injection with Carprofen (5 mg/kg) for post-operative analgesia. For in vivo optogenetics experiments, the AAV5-EF1α-DIO-hChR2(H134R)-mCherry (Addgene plasmid # 37082; Vector Core, University of North Carolina) Control animals received a bilateral injection of a Cre-dependent AAV expressing the cyan fluorescent protein AAV1-EF1α-DIO-CFP (generated in house). For DTA-mediated cell ablation, the AAV1-EF1α-mCherry-flex-dta (Adgene plasmid # 58536 Vector Core, University of North Carolina) was injected bilaterally into the IGL/LGv or the pHB in 3 weeks old *Sox14^Cre^*^/+^ mice. Control animals received a bilateral injection of AAV1-EF1α-DIO-CFP (generated in house).

For monosynaptic tract tracing, we first injected equimolar ratio of AAV1-EF1α-Flex-TVA-mCherry (Addgene plasmid # 38044 Vector Core, University of North Carolina) and AAV1-CMV-DIO-oG (codon optimised; Addgene plasmid # 74290, Vector Core, University of North Carolina) unilaterally in the IGL/LGv or pHB region, followed two weeks later by a second stereotaxic injection of the RV-dG-GFP in the IGL/LGv, SCN or pHB region. Animals were sacrificed 7 days after the RVdG injection.

To identify axonal projections of *Sox14^+^* pHB neurons the AAV1-EF1α-DIO-mGFP (generated in house) or AAV1-phSyn1(S)-FLEX-tdTomato-T2A-SypEGFP-WPRE (Addgene plasmid # 51509, generated in house) were injected unilaterally in the pHB region. Animals were perfused 4 weeks later.

### Brain immunohistochemistry and RNA in situ hybridisation

Mice were transcardially perfused with 4% PFA in PBS and the brains post-fixed at 4 °C overnight. Brains for ISH were stored in PFA for 5 days, to minimise RNA degradation, and all subsequent solutions were treated with diethyl pyrocarbonate (DEPC; AppliChem). The brains were cryoprotected in a sucrose gradient (10–20–30%), frozen on dry ice and cut on a cryostat (Leica) at 60 for IHC on floating sections or cryosectioned at 20μm with coronal sections collected on Superfrost Ultra Plus slides (Thermo Scientific) for ISH.

Immunohistochemistry was performed on floating brain sections. Primary antibodies were incubated on sections twice overnight at 4 °C: chicken anti-GFP (1:10000, Abcam, ab13970), rat anti-RFP (1:1000, 5f8-100, Chromotek), mouse anti-TH (1:1000, MAB5280, Millipore), rabbit anti-calbindin 2 (1:200, ab702, Abcam), rabbit anti-vGAT (1:2000, 131013, Synaptic systems), goat anti-ChAT (1:1000, AB144P, Millipore), mouse anti-TPH (1:50, T0678, Sigma-Aldrich), goat anti-orexin A (1:1000, sc-8070, Santa Cruz Biotechnology), rabbit c-fos (1:800; ABE457 Sigma), goat anti-orexin B (1:1000, sc-8071, Santa Cruz Biotechnology), rabbit anti-MCH (1:1000, H-070-47, Phoenix). Secondary antibodies were incubated on sections for 2 hours at room temperature at a 1:500 dilution. The secondary antibodies used were Alexa-conjugated goat anti-chicken Alexa-488 (A11039, ThermoFisher), goat anti-rat Alexa-568 (A11077, Invitrogen), and goat anti-mouse far red (A11036, Invitrogen), donkey anti-goat Alexa-568 (A11057, ThermoFisher), donkey anti-chicken Alexa-488 (703-545-155, Jackson ImmunoReasearch), donkey anti-mouse far red (ab150107, Millipore), donkey anti-rabbit Alexa-647 (A31573, Invitrogen), donkey anti-rat Alexa-568 (Invitrogen). Blocking and antibody binding solutions where 7% goat serum/PBS with 0.3% TritonX-100 or 3-10% donkey serum/PBS with 1% BSA and O.5% TritonX-100 as blocking solution. After DAPI staining (1:40000 in 1X PBS; Life Technologies) the sections were mounted on Menzel-Glasser Superfrost Plus (J1800AMNZ, ThermoScientific) glass slides using the ProLong Gold antifade reagent (P36930, Invitrogen) mounting medium.

In situ hybridisation (ISH) was performed with a *Npy* antisense RNA probe transcribed *in vitro* from a cDNA template (IMAGE ID: 5683102). The probe was diluted to a final concentration of 800ng/ml in hybridization buffer (50% formamide, 10% dextran sulphate, 1mg/ml rRNA, 1X Denhardt’s solution, 0.2M NaCl, 10mM Tris HCl, 5mM NaH2PO4.2H2O, 1mM Tris base, 50mM EDTA) and applied onto the slides, which were incubated in a humidified chamber at 65°C overnight. The slides were then washed three times for 30min in wash buffer (50% formamide, 1X SSC, 0.1% Tween) at 65°C, two times for 30min in MABT buffer (100mM maleic acid, 150mM NaCl, 0.1% Tween-20) at RT, and blocked for 2h at RT (2% Boehringer Blocking Reagent (Roche), 20% inactivated sheep serum in MABT). Sheep a-DIG alkaline phosphatase conjugated antibody (Roche, 11093274910) was diluted 1:2000 in the blocking solution and incubated with the slides overnight at 4°C. This was followed by five 20min washes in MABT and two 20min washes in the AP buffer (100mM Tris-HCl pH9.5, 100mM NaCl, 50mM MgCl2, 0.1%-Tween-20). NBT/BCIP (Sigma) was diluted in the AP buffer and applied onto the slides for colour reaction for 3–6 hours at RT in the dark.

### Retina immunohistochemistry

The eyes were dissected, post-fixed in 4% PFA overnight, and washed for at least 1 day in PBS at 4°C. The retinas were then dissected in ice cold PBS and washed again in PBS at 4°C. The retinas were then blocked in 10% normal donkey serum, 1% bovine serum albumin (BSA), 0.5% TritonX-100 in PBS for 1h at RT. The following primary antibodies were used: goat anti-ChAT, 1:200 (Chemicon, AB144P); chicken anti-GFP, 1:5000 (Abcam, ab13970); rabbit anti-Opn4, 1:5000 (Advanced Targeting Systems, AB-N38, AB-N39); rabbit anti-UV cone opsin, 1:200 (Millipore, AB5407). The antibodies were diluted in 3% normal donkey serum, 1% BSA, 0.02% sodium azide, 0.5% TritonX-100 in PBS). The retinas were incubated in primary antibodies for 7 days at RT on a shaker. This was followed by three PBS washes, each for 30 min. The secondary antibodies used were donkey anti-goat AlexaFluor 633, 1:500 (ThermoFisher, A21082), donkey anti-goat AlexaFluor 647 (Invitrogen, A21447), donkey anti-rabbit AlexaFluor 568, 1:500 (Invitrogen, A10042) and donkey anti-chicken AlexaFluor 488, 1:500 (Jackson ImmunoReasearch, 703-545-155), diluted in 3% NDS, and incubated with the retinas for 1 day at 4°C. The next day, there were two 30 min PBS washes, followed by incubation in DAPI (1:40000 in PBS; Life Technologies) overnight at 4°C. The retinas were then changed to PBS and mounted, using the ProLong Diamond mounting medium (Invitrogen). Spacers (SLS, 24 × 24 mm, no.15) were used on the slides to prevent the coverslips compressing the retinas.

### Optogenetic stimulation

Animals were chronically tethered to a branching fibreoptic patch cord (200 µm diameter core, 0.53 NA; Doric Lenses) attached to the implanted cannula and connected to a high-powered blue (470 nm) LED (Doric Lenses) under the control of an LED Driver (LEDRVP-2CH, Doric Lenses). LED source and patch cord were connected via an optical rotary joint allowing free movements of the animal in a circular cage. Mice were kept in constant darkness and allowed to free run at least a week before stimulation. Locomotor activity was monitored in 1 min bins using Clocklab software (Actimetrics, Inc, Wilmette, IL, USA). Light pulses (470 nm, 8 Hz, 10 ms duration, 1h) were generated approximately 3 hours or 6 hours after the onset of the active phase, through Doric Neuroscience Studio software (Doric Lenses) and repeated daily at the same clock time over 14 days. Light intensity at the cannula tip was determined to be 4.5mV when driven at 1000 mA using a PM100D Optical Power Meter (Thorlabs).

### Light exposure protocol

Mice were single-housed in a circadian light-, air-, temperature-controlled ventilated cabinet (Phenome Technologies) monitored by Clocklab Chamber Control Software (Actimetrics, Inc, Wilmette, IL, USA). Mice were first entrained to 12 h: 12 h light dark cycle under standard light intensity (200 lux). Then, all subjects went through a “jet-lag” paradigm (6 h phase advance) using bright light (200 lux) lasting 14 days. Mice were then housed in constant darkness for 14 days and free running was assessed. Following these light conditions, mice were allowed to re-entrained to 12 h: 12 h light dark cycle under bright light (200 lux) for 2 weeks before going through a novel “jet-lag” paradigm (6 h phase advance) using dim light (10 lux) lasting 14 days. General activity was measured by using infrared motion sensors (Actimetrics, Inc, Wilmette, IL, USA) wired to a computer. Data were collected in 1-min bins using Clocklab software (Actimetrics, Inc, Wilmette, IL, USA).

### Behavioural tests

Sucrose anhedonia. Adult mice (approx. 9 months old) were single housed in the presence of two water bottles 1 day before testing to acclimate them to the bottles. Sucrose preference was assessed over 2 days. On the first day, one bottle containing 1% sucrose and one bottle containing water were introduced at the beginning of the active phase (7pm). Bottles were removed at the end of the active phase (7am). On the second day, the position of the two bottles was switched and the procedure repeated. Bottles were weighed at the beginning and end of the active period to measure amount consumed as well as mice and food to detect any abnormal change in food consumption. Sucrose preference was calculated by dividing the amount of sucrose consumed by the total amount consumed (water and sucrose). The percentage of sucrose consumed by control and ablated mice was compared by the Mann Whitney *U* test.

Forced swim test. Adult mice (approx. 9 months old) were individually placed in an inescapable clear Perspex cylinder (49 cm high x 15 cm diameter), filled with 40 cm of water at 25°C. Mice were carefully placed in the water and left to swim for 6 min. Behaviour was monitored by a video camera positioned in front of the apparatus and scored manually through Ethovision software (Noldus). Time spent immobile for the last 4 min of the test was calculated. Increased time spent immobile is indicative of increased depression-related behaviour. The amount of time spent immobile during the last 4 min was analysed by the Mann Whitney *U* test.between control and ablated mice.

Elevated-plus maze. The apparatus consisted of two open arms (30 × 5 cm) opposite to one another and two arms enclosed by opaque walls (30 × 5 × 15 cm) opposite of one another forming a cross. The arms were separated by a central platform (5 × 5 cm). The maze was elevated (40 cm) such that the open arms are aversive due to openness, unfamiliarity and elevation. The light intensity in the open arms was 200 lux, whereas the light intensity in the closed arms was 10 lux. Mice were placed in the centre of the elevated plus maze facing one of the open arms and allowed to freely move around the maze for 5 min. Behaviour was monitored from above by a video camera connected to Ethovison video tracking system (Noldus). The apparatus was cleaned thoroughly between each trial. The time spent and the distance travelled in the open arms were measured as indications of anxiety-related behaviour. These measures were compared between control and ablated mice using Student’s unpaired t-test.

### Sleep recording

Adult mice (approx. 6 months old) were chronically implanted with screw-type electrodes in the skull to measure cortical EEG. A pair of stainless-steel electrodes was implanted in the dorsal neck muscle to measure EMG. Screw electrodes were placed in burr holes in the skull over the parietal cortex (–1.5 mm Bregma, +1.5 mm midline) and frontal cortex (+1.5 mm Bregma, –1.5 mm midline) with a reference electrode over the cerebellum (1.0 mm caudal to lambda, 0 mm midline) and a ground over the olfactory bulb area.

Electrodes were connected to head-mounts and secured with dental cement. The animals were allowed to recover from surgery for at least one week, before the EEG/EMG recordings were performed.

At the time of the recordings, mice were tethered to four channel EEG/EMG recording systems (Pinnacle Technology Inc.) and housed individually and sequentially in a soundproof and light-controlled cabinet (standard light conditions 200 lux) equipped with a videocamera with a 3.6 mm lens and infrared illumination (Pinnacle Technology Inc). Data was acquired continuously for a 48-hour period, maintaining the same light-dark cycle, temperature and humidity as for the home cages. The EEG/EMG signals were sampled at 250 Hz, amplified 100× and low-pass filtered at 100 Hz using a two EEG channel, two EMG channel mouse pre-amplifier (Pinnacle Technology Inc).

Sleep scoring was performed manually on 10-second epochs using Sirenia Sleep software (Pinnacle Technology Inc.). EEG and EMG recordings were synchronized for each epoch to video recordings. Epochs with EMG amplitude slightly (quiet WAKE) or significantly higher than baseline (active WAKE), together with desynchronized low amplitude EEG were scored as “WAKE”. Epochs with low-amplitude EMG and high amplitude delta (1-4 Hz) activity were scored as “NREM” and epochs with low amplitude EMG accompanied by low-amplitude rhythmic theta activity (6-9 Hz) were recorded as “REM” (Quattrocchi et al., 2015).

Distance travelled and velocity were extracted from video files at 30 fps and synchronized with the EEG and EMG data for each individual mouse, using the Sirenia software video plugin (Pinnacle Technology, Inc).

### Electrophysiology on acute brain slices

Animals were culled in accordance with the UK Home Office guidelines. Brains were rapidly removed from the skull and immediately immersed in ice cold slicing solution (92 mM NMDG, 2.5 mM KCl, 1.25 mM NaH2PO4, 30 mM NaHCO3, 20 mM HEPES, 25 mM glucose, 2 mM thiourea, 5 mM Na-ascorbate, 3 mM Na-pyruvate, 0.5 mM CaCl2·4H2O and 10 mM MgSO4·7H2O), pH 7.3-7.4 when bubbled with 95%O2/5%CO2). Slices were cut using a vibratome tissue slicer (Campden instruments) at a thickness of 300 μm, after which they were immediately transferred to a holding chamber containing slicing NMDG at 33-34°C continuously bubbled with 95%O2/5%CO2. Slices were left to equilibrate for 10-15 minutes, after which they were transferred into a holding chamber at room temperature containing recording ACSF (125 mM NaCl, 2.5 mM KCl, CaCl2 2 mM, 1 mM MgCl, 1.25 mM NaH2PO4, 26 mM NaHCO3, 11 mM glucose, pH 7.4) that was continuously bubbled with 95%O2/5%CO2. Slices were then visualized using a fixed-stage upright microscope (BX51W1, Olympus and Scientifica Slice scope) fitted with a high numerical aperture water-immersion objective and an infra-red sensitive digital camera. A 595nm amber LED was used for identifying mCherry expression and a 470nm blue LED was used for optogenetics stimulation. Patch pipettes were made from thick-walled borosilicate glass capillaries (0.86 mm internal diameter, 1.5 mm outer diameter, Harvard Apparatus) using a two-step vertical puller (Narishige, PC-10). Pipette resistances were typically 5-8 MΩ when back filled with internal solution. For voltage-clamp experiments, the internal solution contained: 140 mM CsCl, 4 NaCl mM, 0.5 mM CaCl2, 10 mM HEPES, 5 mM EGTA, 2 Mg-ATP mM; and the pH was adjusted to 7.3 with CsOH. For current-clamp experiments the internal solution contained: 145 mM K-gluconate; 4 mM NaCl; 0.5 mM CaCl2; 10 mM HEPES; 5 mM EGTA; 4 mM Mg-ATP; 0.3 mM Na-GTP (adjusted to pH 7.3 with KOH). The amplifier head stage was connected to an Axopatch 700B amplifier (Molecular Devices; Foster City, CA).

The amplifier current output was filtered at 10 kHz (–3 dB, 8-pole low-pass Bessel) and digitized at 20 kHz using a National Instruments digitization board (NI-DAQmx, PCI-6052E; National Instruments, Austin, Texas). Data acquisition was performed using CED Signal (Version 6) software. CED Signal’s “IntraSpikeAnalysis” spike detection script was used for Current Clamp action potential detection thresholding at 0 mV. WinEDR (Strathclyde Electrophysiology Software) was used for Voltage Clamp postsynaptic current detection through template fitting at 0.1ms (Tau Rise) & 10ms (Tau Decay).

For Peristimulus Time Histogram (PSTH), an in-house MATLAB code (https://github.com/dd119-ic/BrockManuscript) was used to construct PSTHs from the optogenetic input timings and detected events. OriginPro v2020 was used to construct histograms and for power spectrum analysis.

## QUANTIFICATION AND STATISTICAL ANALYSIS

### Monosynaptic viral tracing

Nikon A1R Inverted or Nikon Upright Ni-E confocal optics were used to acquire images using a 20X/NA 0.75 Plan Apo VC or a 60X/1.4NA objective. Data on starter cells were collected by analysing z-stack images of all coronal sections spanning the entire injection site, acquired with A1R Nikon confocal microscopes, using the ‘multipoint’ function in Fiji (Schindelin et al., 2012). Mono-synaptic inputs were calculated as percent of total for each brain or normalised per starter cell, using Excel 365 (Microsoft) and GraphPad Prism 8 software.

The location and distribution of transynaptically labelled neurons across the brain was assessed using a Zeiss AxioImager microscope using the 4X/0.10 Acroplan and 10X/0.3 Ph1 EC-Plan-NeoFluar objectives. Regions where the GFP^+^ somas were present were identified by comparison with the Paxinos & Franklin Mouse Brain Atlas. The Zeiss AxioImager microscope was also used to acquire overviews of coronal sections showing the brain-wide distribution of inputs, using a Plan NeoFluar 2.5X/0,075 objective.

### RGC analysis

A Nikon A1R confocal microscope was used to acquire z-stacks (step size 1.1μm) of the RGCs using the 20X/NA 0.75 Plan Apo VC objective. The stacks were acquired such that both ON and OFF ChAT^+^ layers and the entire extent of the RGC’s dendrites were included. Overviews of the retinas were acquired using a 10X/NA 0.3 Plan Fluor D objective and images were acquired as z-stacks (step size: 10 μm) and were composed of 4×4 tiles.

RGC dendritic arbour tracing and annotation of ChAT layers was based on the protocol published in (Sumbul et al., 2014). The RGC dendritic trees were traced manually using the Simple Neurite Tracer plugin (Longair et al., 2011) and SNT plugin in Fiji (Schindelin et al., 2012) and exported as .swc files. To annotate the ChAT layers, the z-stack images were first resliced so that the z-dimension was projected onto the y-axis. The ON ChAT layer was manually annotated, using the ‘multipoint’ function in Fiji, and every 80th digital slice was marked with 5–10 data points. The x, y and z coordinates for all the points were exported as a .txt file. The procedure was then repeated for the OFF ChAT layer. Opn4 expression levels in RVdG-infected RGCs was compared with the stronger signal from putative M1 ipRGCs and the background signal in the RGC layer, within the same image frame.

RVdG-labelled Opn4^+^ RGCs were analysed using a MATLAB implementation of the algorithm developed by (Sumbul et al., 2014) and available at https://github.com/padraic-padraic/rgc. The algorithm begins by ‘unwarping’ the ChAT layers, to produce two flat planes corresponding to the ON and OFF layers. The program then quantifies the arbour density in the IPL relative to these layers, outputting a histogram of arbour density against the IPL z-axis, where z=0μm corresponds to the ON layer and z=12μm corresponds to the OFF layer. The stratification data was then processed further, based on (Siegert et al., 2009) and (Rompani et al., 2017). In particular, the IPL was divided into 10 layers, which were defined such that the OFF ChAT layer is contained in strata 3, and the ON layer in strata 7. The remaining layers were defined by linearly interpolating the spacing between the ON and OFF ChAT layers and extending this interpolation to produce 10 full strata corresponding to 3 μm each. The dendritic arbour density histogram was then binned into each of these 10 strata using our custom MATLAB script (https://github.com/padraic-padraic/rgc). Each bin contained the summed arbour density within its range. Stratification above and below the ‘boxed’ region was included in bins 1 and 10, respectively. A stratum was considered ‘labelled’ if the total arbour density in that bin was greater than the average density across all bins. This strata labelling was used to output boxplots of stratification. Cells were then classified into ON, OFF or ON-OFF stratifying using the labelled stratum. ON stratifying cells had only stratum in bins 6-10 labelled. OFF stratifying cells had only stratum bins 1-5 labelled. Lastly, ON-OFF stratifying cells had either stratum labelled from stratum bins 1-5 and 6-10 or were seen to have clear peaks in ranges corresponding to both ON and OFF stratum bins from raw stratification output.

The diameter of the dendritic arbour was quantified using our custom MATLAB script (https://github.com/urygarsumbul/rgc), based on the skeletonized arbour. The output .swc file contains a description of the arbour as ‘nodes’ connected by ‘edges’. The ends of the dendritic arbor were identified as all nodes connected to only a single edge. Their x and y co-ordinates were converted to physical distances from the centre of the image by multiplying them with the corresponding voxel resolutions in μm. Using the MATLAB ‘pdist’ routine, the Euclidian distance between every pair of end-nodes was calculated, and the maximum value was taken as the dendritic diameter. Scholl analysis and measurements of total branching points and total dendritic length were performed using the SNT plugin for FIJI. Total branching points and total dendritic length were measured using SNTs in-built measurement functions for cable length and number of branch points. In Scholl analysis a pixel central to the nucleus of RGCs was chosen as a starting point and radius step size was set to 10 µm. Statistical comparisons of morphological parameters from ON, OFF and ON-OFF stratifying cell groups were made with Kruskal-Wallis tests followed by Dunn’s multiple comparisons tests.

### Behavioural data analysis

Investigators were blinded to the group allocation during experiments or data analysis. Sample sizes were indicated in the figures and associated text and are similar to those reported in previous studies (An et al., 2020; Huang et al., 2019). Data were analysed using the GraphPad Prism 8 software. For all statistical comparisons, we first analysed the data distribution with the Shapiro–Wilk test and D’Agostino-Pearson test for normality. Statistical differences of normally distributed data were then determined using unpaired two-tailed *t*-test. The Mann-Whitney *U* test was employed to analyse non-normally distributed data. *P* values less than 0.05 were considered significant and the thresholds for *P* value significance were reported as follows: * *P* < 0.05; \*\**P* ≤ 0.01; \*\*\**P* ≤ 0.001; \*\*\*\**P* ≤ 0.0001. Experimental errors are presented as standard error of the mean (s.e.m.).

### EEG/EMG Data Analysis

Cumulative power in alpha (8-12 Hz), delta (0.5-4 Hz) and theta (6-9 Hz) frequency bands were calculated by computing the discrete Fourier transform (DFT) of the EEG data using a fast Fourier transform (FFT) algorithm. Then, the summed power in each frequency band was normalised to the sum of the power over the entire range (0–15 Hz). Power Spectral Density (PSD) estimates were calculated using Welch method (window length = 1000; NFFT = 1024). Spectograms showing the amplitude of EEG signals in the time and frequency domain were generated using Short-time Fourier transform (window length = 1024; NFFT = 4096), as previously described (Zhivomirov, 2019). Theta/Delta and Delta/Theta ratios were calculated by summing PSD values for each frequency range and dividing by each other. EEG signal analyses were conducted using custom codes written in MATLAB, as previously described (Gelegen et al., 2014; Gelegen et al., 2018; Pang et al., 2009; Zhivomirov, 2019).

EEG/EMG statistical analysis: All statistical tests were performed in GraphPad Prism 8. Shapiro-Wilk test was used for normality of distribution. Data are represented as the mean ± s.e.m., unless otherwise stated. Time spent at each vigilance state, Theta/Delta and Delta/Theta ratios for the hour preceding and following the circadian light change was compared between the two groups using paired t-test or Wilcoxon test, depending on the normality of the data. Normalized power at Alpha, Delta and Theta frequency bands for the hour preceding and two hours following the circadian light change were compared between the groups first using repeated measures One Way ANOVA or Friedman Test depending on the normality of the data. If a significant overall p value was obtained, individual time points were compared using paired t-test or Wilcoxon test.

### Code availability

The code for RGC analysis generated during this study is available at GitHub https://github.com/padraic-padraic/rgc.

The code for Peristimulus Time Histogram (PSTH) is available at GitHub

https://github.com/dd119-ic/BrockManuscript

## Acknowledgments

We are grateful to Nicola Maiorano, Kamill Balint, Botond Roska at the Friedrich Miescher Institute (FMI, CH) and Marco Tripodi MRC Laboratory of Molecular Biology (LMB, University of Cambridge, UK) for reagents and protocols for recombinant rabies virus production. We are grateful to Samer Hattar (NIH, US) for the gift of the Opn4taulacZ mouse line. We thank Padraic Calpin (University College London, UK) for designing the MATLAB code to analyse RGCs. We thank Vladyslav Vyazovskiy (University of Oxford, UK) for advice on EEG/EMG recordings in mice. We thank the Wohl Cellular Imaging Centre (WCIC) at Kings College London for help with microscopy. This work was funded by a Royal Society grant RG160741 and Biotechnology and Biological Sciences Research Council (BBSRC) BB/L020068/1 and BB/R007020/1 grants to AD. OB was the recipient of an Independent Research Award by the Institute of Psychiatry, Psychology and Neuroscience, King’s College London.

## Extended figure and tables

**Table 4-1.**
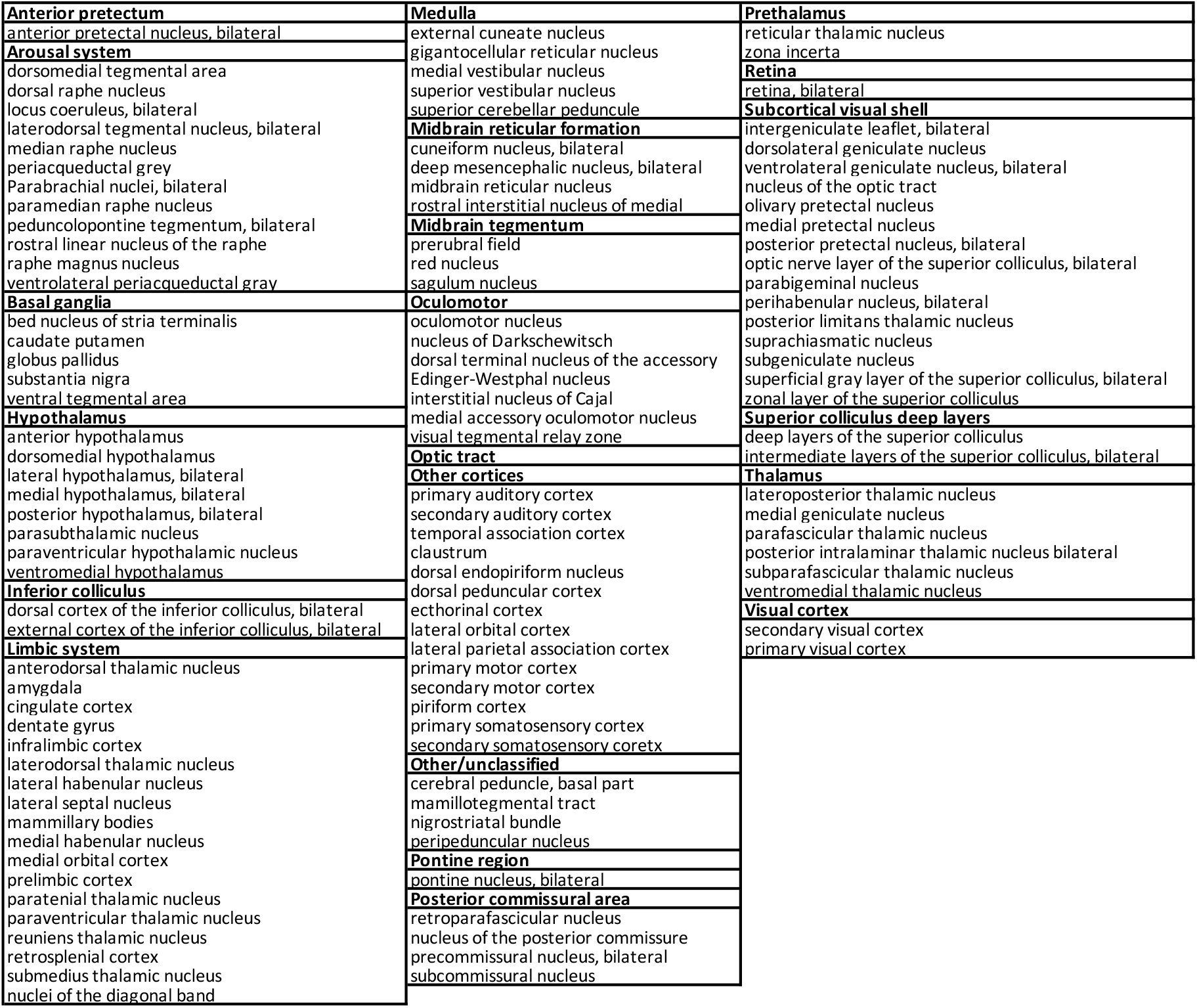
Anatomical classification of regions harbouring presynaptic input to *Sox14^+^* neurons. The classification used to cluster anatomical regions containing cells transsynaptically labelled by the RVdG vector is based on the atlas of the mouse brain by Paxinos and Franklin (Paxinos and Franklin, 2001).

**Table 6-1.** Statistical treatment of EEG/EMG data. Mean values, experimental error and the parametric and non-parametric tests used to calculate statistical significance. F indicates Friedman test; W indicates Wilcoxon.

| Parameter | Mean ± s.e.m. | t / Wilcoxon (W) statistics | RM ANOVA / Friedman (F) statistics | p value | Figure # |
| --- | --- | --- | --- | --- | --- |
| **24 hours period** |  |  |  |  |  |
| Total Wake (Control vs Ablated) | 48.24 ± 1.96 vs 48.22 ± 1.71 | 0.0072 | N/A | 0.99 | 4C |
| Total NREM (Control vs Ablated) | 46.73 ± 1.80 vs 46.38 ± 1.71 | 0.139 | N/A | 0.89 | 4C |
| Total REM (Control vs Ablated) | 5.03 ± 0.20 vs 5.39 ± 0.17 | 1.326 | N/A | 0.2 | 4C |
| **2 hours before and 2 hours after the light to dark transition** |  |  |  |  |  |
| Wake L/D Control | N/A | N/A | 24.2 | **<0.0001** | 4D |
| Wake L/D Ablated | N/A | N/A | 6.444 | **0.015** | 4D |
| NREM L/D Control | N/A | N/A | 22.46 | **0.0001** | 4D |
| NREM L/D Ablated | N/A | N/A | 6.092 | **0.018** | 4D |
| REM L/D Control | N/A | N/A | 12.45 (F) | **0.0005** | 4D |
| REM L/D Ablated | N/A | N/A | 4.66 (F) | 0.099 | 4D |
| **2 hours before and 2 hours after the dark to light transition** |  |  |  |  |  |
| Wake D/L Control | N/A | N/A | 14 (F) | **<0.0001** | 4E |
| Wake D/L Ablated | N/A | N/A | 5.85 (F) | 0.051 | 4E |
| NREM D/L Control | N/A | N/A | 14 (F) | **<0.0001** | 4E |
| NREM D/L Ablated | N/A | N/A | 4.84 (F) | 0.09 | 4E |
| REM D/L Control | N/A | N/A | 12.25 (F) | **0.0003** | 4E |
| REM D/L Ablated | N/A | N/A | 8.27 (F) | **0.019** | 4E |
| **1 hour before compared to the first or second hour after the light to dark transition** |  |  |  |  |  |
| Wake_Control_L(-1) vs D(+1) | 33.17 ± 2.98 vs 80.26 ± 3.49 | 8.066 | N/A | **<0.0001** | 4F |
| Wake_Control_L(-1) vs D(+2) | 33.17 ± 2.98 vs 72.61 ± 7.76 | 5.366 | N/A | **0.001** | 4F |
| Wake_Ablated_L(-1) vs D(+1) | 36.57 ± 5.34 vs 56.67 ± 3.49 | 1.992 | N/A | 0.093 | 4F |
| Wake_Ablated_L(-1) vs D(+2) | 36.57 ± 5.34 vs 75.23 ± 7.24 | 3.899 | N/A | **0.008** | 4F |
| NREM_Control_L(-1) vs D(+1) | 60.08 ± 2.65 vs 18.39 ± 3.29 | 7.768 | N/A | **0.0001** | 4F |
| NREM_Control_L(-1) vs D(+2) | 60.08 ± 2.65 vs 25.17 ± 7.04 | 5.051 | N/A | **0.0015** | 4F |
| NREM_Ablated_L(-1) vs D(+1) | 56.35 ± 4.73 vs 38.19 ± 7.68 | 1.992 | N/A | 0.093 | 4F |
| NREM_Ablated_L(-1) vs D(+2) | 56.35 ± 4.73 vs 22.36 ± 6.64 | 3.89 | N/A | **0.0081** | 4F |
| REM_Control_L(-1) vs D(+1) | 6.75 ± 0.52 vs 1.35 ± 0.45 | 6.363 | N/A | **0.0004** | 4F |
| REM_Control_L(-1) vs D(+2) | 6.75 ± 0.52 vs 2.18 ± 0.85 | -36 (W) | N/A | **0.007** | 4F |
| REM_Ablated_L(-1) vs D(+1) | 7.08 ± 0.78 vs 5.10 ± 1.16 | -14 (W) | N/A | 0.296 | 4F |
| REM_Ablated_L(-1) vs D(+2) | 7.08 ± 0.78 vs 2.41 ± 0.90 | -26 (W) | N/A | **0.031** | 4F |
| **1 hour before compared to the first or second hour after the dark to light transition** |  |  |  |  |  |
| Wake_Control_D(-1) vs L(+1) | 94.77 ± 3.48 vs 63.47 ± 9.42 | -28 (W) | N/A | **0.015** | 4G |
| Wake_Control_D(-1) vs L(+2) | 94.77 ± 3.48 vs 36.11 ± 9.72 | -28 (W) | N/A | **0.015** | 4G |
| Wake_Ablated_D(-1) vs L(+1) | 79.18 ± 7.51 vs 82.47 ± 7.80 | 3 (W) | N/A | 0.812 | 4G |
| Wake_Ablated_D(-1) vs L(+2) | 79.18 ± 7.51 vs 39.37 ± 8.09 | 3.254 | N/A | **0.017** | 4G |
| NREM_Control_D(-1) vs L(+1) | 5.22 ± 3.48 vs 34.87 ± 8.82 | 28 (W) | N/A | **0.015** | 4G |
| NREM_Control_D(-1) vs L(+2) | 5.22 ± 3.48 vs 57.51 ± 8.86 | 28 (W) | N/A | **0.015** | 4G |
| NREM_Ablated_D(-1) vs L(+1) | 20.50 ± 7.34 vs 17.25 ± 7.65 | -3 (W) | N/A | 0.848 | 4G |
| NREM_Ablated_D(-1) vs L(+2) | 20.50 ± 7.34 vs 56.31 ± 7.22 | 3.078 | N/A | **0.021** | 4G |
| REM_Control_D(-1) vs L(+1) | 0.00 ± 0.00 vs 1.66 ± 0.64 | 10 (W) | N/A | 0.125 | 4G |
| REM_Control_D(-1) vs L(+2) | 0.00 ± 0.00 vs 6.37 ± 1.09 | 28 (W) | N/A | **0.015** | 4G |
| REM_Ablated_D(-1) vs L(+1) | 0.31 ± 0.21 vs 0.27 ± 0.20 | -1 (W) | N/A | >0.999 | 4G |
| REM_Ablated_D(-1) vs L(+2) | 0.31 ± 0.21 vs 4.31 ± 1.34 | 26 (W) | N/A | **0.0313** | 4G |
| **24 hours period** |  |  |  |  |  |
| Total Distance_12hr_D vs L_Control | 100 ± 14.54 vs 77.49 ± 26.22 | -18 (W) | N/A | 0.25 | 4H |
| Total Distance_12hr_D vs L_Ablated | 100 ± 17.38 vs 51.82 ± 8.19 | -26 (W) | N/A | **0.031** | 4H |
| **1 hour before and 1 hour after the light to dark transition** |  |  |  |  |  |
| Total Distance_One Hour_L vs D_Control | 100 ± 21.26 vs 559.3 ± 136.2 | 36 (W) | N/A | **0.0078** | 4I |
| Total Distance_One Hour_L vs D_Ablated | 100 ± 16.65 vs 279.9 ± 67.93 | 2.539 | N/A | **0.044** | 4I |
| **1 hour before and 1 hour after the dark to light transition** |  |  |  |  |  |
| Total Distance_One Hour_D vs L_Control | 100 ± 15.13 vs 64.49 ± 31.44 | -22 (W) | N/A | 0.148 | 4J |
| Total Distance_One Hour_D vs L_Ablated | 100 ± 28.52 vs 92.37 ± 29.29 | -10 (W) | N/A | 0.468 | 4J |

**Table7-1.** Statistical treatment of EEG/EMG data. Mean values, experimental error and the parametric and non-parametric tests used to calculate statistical significance. F indicates Friedman test; W indicates Wilcoxon.

| Parameter | Mean ± s.e.m. | t / Wilcoxon (W) statistics | RM ANOVA / Friedman (F) statistics | p value | Figure # |
| --- | --- | --- | --- | --- | --- |
| **1 hour before and 1 hour after the light to dark transition** |  |  |  |  |  |
| Theta/Delta_Control_L(-1) vs D(+1) | 0.87 ± 0.01 vs 0.97 ± 0.03 | 4.996 | N/A | **0.001** | 5B |
| Theta/Delta_Ablated_L(-1) vs D(+1) | 0.86 ± 0.01 vs 0.89 ± 0.01 | 22 (W) | N/A | 0.078 | 5E |
| **1 hour before and 1 hour after the dark to light transition** |  |  |  |  |  |
| Theta/Delta_Control_D(-1) vs L(+1) | 0.98 ± 0.02 vs 1.08 ± 0.04 | 3.123 | N/A | **0.016** | 5H |
| Theta/Delta_Ablated_D(-1) vs L(+1) | 1.06 ± 0.05 vs 1.07 ± 0.05 | 0.517 | N/A | 0.623 | 5K |
| **1 hour before compared to the first or second hour after the light to dark transition** |  |  |  |  |  |
| Delta_Control_Light/Dark | N/A | N/A | 7.672 | **0.006** | 5M |
| Delta_Ablated_Light/Dark | N/A | N/A | 0.857(F) | 0.768 | 5M |
| Delta_Control_L(-1) vs D(+1) | 0.29 ± 0.009 vs 0.26 ± 0.01 | 3.509 | N/A | **0.009** | 5M |
| Delta_Control_L(-1) vs D(+2) | 0.29 ± 0.009 vs 0.28 ± 0.007 | 1.063 | N/A | 0.323 | 5M |
| Delta_Ablated_L(-1) vs D(+1) | 0.30 ± 0.007 vs 0.30 ± 0.009 | -5(W) | N/A | 0.734 | 5M |
| Delta_Ablated_L(-1) vs D(+2) | 0.30 ± 0.007 vs 0.29 ± 0.013 | -2(W) | N/A | 0.937 | 5M |
| **1 hour before compared to the first or second hour after the light to dark transition** |  |  |  |  |  |
| Theta_Control_Light/Dark | N/A | N/A | 12.02 | **0.001** | 5N |
| Theta_Ablated_Light/Dark | N/A | N/A | 2(F) | 0.486 | 5N |
| Theta_Control_L(-1) vs D(+1) | 0.23 ± 0.003 vs 0.27 ± 0.009 | 4.535 | N/A | **0.002** | 5N |
| Theta_Control_L(-1) vs D(+2) | 0.23 ± 0.003 vs 0.25 ± 0.006 | 3.2 | N/A | **0.105** | 5N |
| Theta_Ablated_L(-1) vs D(+1) | 0.24 ± 0.003 vs 0.25 ± 0.007 | 20(W) | N/A | 0.109 | 5N |
| Theta_Ablated_L(-1) vs D(+2) | 0.24 ± 0.003 vs 0.25 ± 0.01 | 20(W) | N/A | 0.109 | 5N |
| **1 hour before compared to the first or second hour after the light to dark transition** |  |  |  |  |  |
| Alpha_Control_Light/Dark | N/A | N/A | 13.31 | **<0.0001** | 5O |
| Alpha_Ablated_Light/Dark | N/A | N/A | 2(F) | 0.486 | 5O |
| Alpha_Control_L(-1) vs D(+1) | 0.24 ± 0.004 vs 0.25 ± 0.005 | 4.231 | N/A | **0.003** | 5O |
| Alpha_Control_L(-1) vs D(+2) | 0.24 ± 0.004 vs 0.24 ± 0.003 | 2.429 | N/A | **0.045** | 5O |
| Alpha_Ablated_L(-1) vs D(+1) | 0.24 ± 0.003 vs 0.24 ± 0.003 | 20(W) | N/A | 0.109 | 5O |
| Alpha_Ablated_L(-1) vs D(+2) | 0.24 ± 0.003 vs 0.25 ± 0.008 | 1.964 | N/A | 0.097 | 5O |
| **1 hour before compared to the first or second hour after the dark to light transition** |  |  |  |  |  |
| Delta_Control_Dark/Light | N/A | N/A | 8.42 | **0.011** | 5P |
| Delta_Ablated_Dark/Light | N/A | N/A | 6(F) | 0.051 | 5P |
| Delta_Control_D(-1) vs L(+1) | 0.24 ± 0.009 vs 0.28 ± 0.016 | 3.036 | N/A | **0.019** | 5P |
| Delta_Control_D(-1) vs L(+2) | 0.24 ± 0.009 vs 0.29 ± 0.016 | 3.13 | N/A | **0.016** | 5P |
| Delta_Ablated_D(-1) vs L(+1) | 0.29 ± 0.017 vs 0.30 ± 0.017 | 14(W) | N/A | 0.296 | 5P |
| Delta_Ablated_D(-1) vs L(+2) | 0.29 ± 0.017 vs 0.32 ± 0.009 | 2.684 | N/A | **0.036** | 5P |
| **1 hour before compared to the first or second hour after the dark to light transition** |  |  |  |  |  |
| Theta_Control_Dark/Light | N/A | N/A | 7(F) | **0.03** | 5Q |
| Theta_Ablated_Dark/Light | N/A | N/A | 5.894 | **0.028** | 5Q |
| Theta_Control_D(-1) vs L(+1) | 0.29 ± 0.011 vs 0.26 ± 0.016 | -32(W) | N/A | **0.023** | 5Q |
| Theta_Control_D(-1) vs L(+2) | 0.29 ± 0.011 vs 0.24 ± 0.014 | -32(W) | N/A | **0.023** | 5Q |
| Theta_Ablated_D(-1) vs L(+1) | 0.26 ± 0.012 vs 0.26 ± 0.012 | 0.396 | N/A | 0.706 | 5Q |
| Theta_Ablated_D(-1) vs L(+2) | 0.26 ± 0.012 vs 0.23 ± 0.003 | 2.648 | N/A | **0.038** | 5Q |
| **1 hour before compared to the first or second hour after the dark to light transition** |  |  |  |  |  |
| Alpha_Control_Dark/Light | N/A | N/A | 12.01 | **0.002** | 5R |
| Alpha_Ablated_Dark/Light | N/A | N/A | 4.571(F) | 0.111 | 5R |
| Alpha_Control_D(-1) vs L(+1) | 0.27 ± 0.005 vs 0.24 ± 0.007 | 3.344 | N/A | **0.012** | 5R |
| Alpha_Control_D(-1) vs L(+2) | 0.27 ± 0.005 vs 0.23 ± 0.006 | 3.975 | N/A | **0.005** | 5R |
| Alpha_Ablated_D(-1) vs L(+1) | 0.26 ± 0.014 vs 0.26 ± 0.014 | -12(W) | N/A | 0.375 | 5R |
| Alpha_Ablated_D(-1) vs L(+2) | 0.26 ± 0.014 vs 0.23 ± 0.004 | -24(W) | N/A | 0.046 | 5R |

**Figure 9-1.**
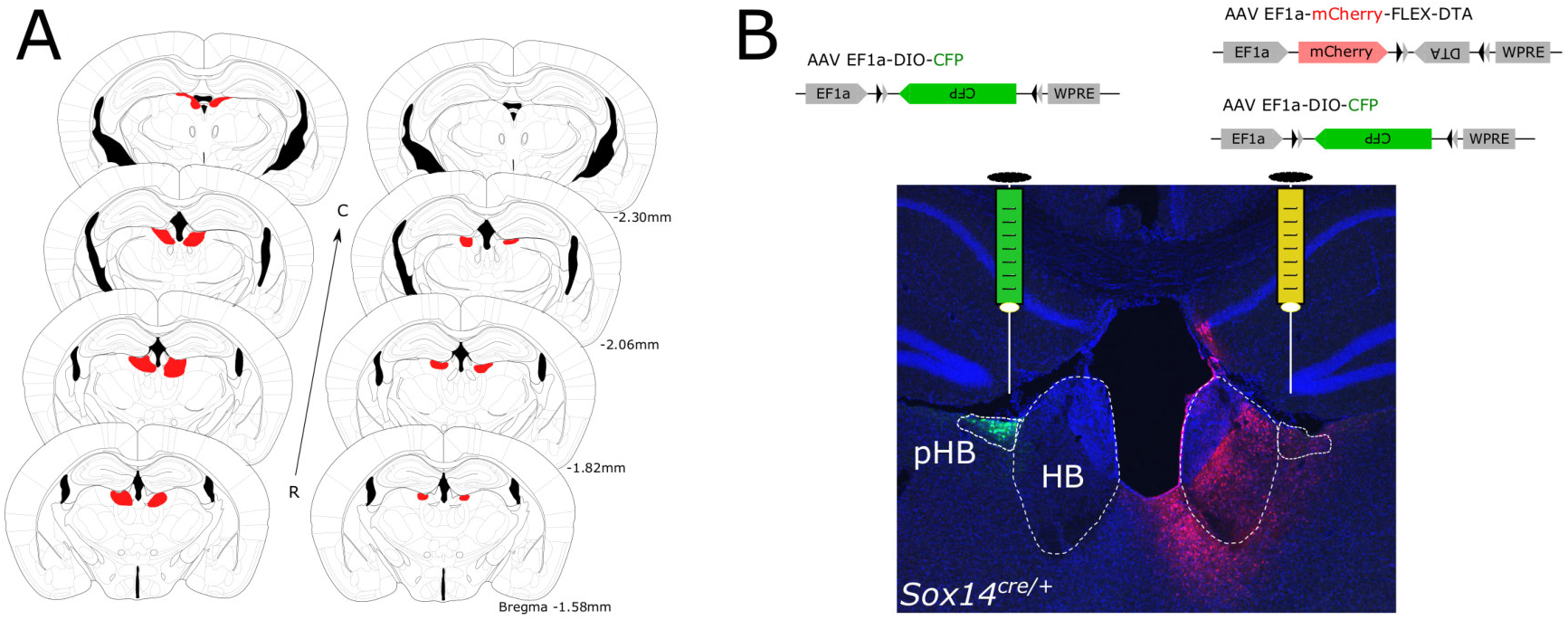
Evaluation of the extent of *Sox14^+^* pHB cell ablation**. *A***, Evaluation of the AAV1-EF1α-mCherry-flex-dta spread upon stereotaxic injection in the pHB region of *Sox14^Cre/+^* mice. Two representative brains depicting the variability and location of mCherry labelled cells (red) in a larger injection (left) and smaller injection (right). ***B***, Illustrative example of the strategy used ablate *Sox14^+^* neurons in the pHB. The AAV1-EF1α-DIO-CFP Cre-dependent AAV vector (green) was injected stereotaxically in the left pHB. An equimolar mixture of the EF1α-DIO-CFP and the EF1α-mCherry-flex-dta was injected in the right pHB. Lack of CFP signal on the right pHB indicate efficient ablation of all infected cells.

